# Conserved catalytic activity of animal TIR domains functions in TLR4 signaling

**DOI:** 10.64898/2026.04.20.719644

**Authors:** Bohdana Hurieva, Sarah B. Christ, Sophia J. Schwojer, Ilya Osterman, Anna-Lena Bergmaier, Alla H. Falkovich, Erez Yirmiya, Azita Leavitt, Jeremy Garb, Ohad Roth, Gil Amitai, Veit Hornung, Rotem Sorek

## Abstract

The Toll/interleukin-1 receptor (TIR) domain is important for immune signaling across bacteria, plants, and animals. In animal innate immunity, TIR domains are thought to function as adaptors mediating protein-protein interactions, yet studies in bacteria and plants revealed that TIR domains often act as enzymes that produce immune signaling molecules. Here, we show that catalytic activity is a conserved property of TIR domains across the animal tree of life. We find that TIR domains from evolutionarily diverse animals, including the TIR domain of human TLR4, are enzymes that can produce cyclic ADP-ribose (cADPR). We further show that the catalytic residue of TLR4 TIR domain is required for TLR4 signal transduction in human macrophages. Substitution of the catalytic glutamate abolishes LPS-induced inflammatory responses and disrupts Myddosome assembly, revealing a previously unrecognized functional requirement for the TIR catalytic site in TLR4 signaling. Beyond animals, we identify a TIR domain from green algae that produces 3ʹcADPR, suggesting deep evolutionary conservation of TIR-mediated signaling. Together, our findings establish enzymatic activity as a widespread and evolutionarily conserved property of TIR domains and reveal a functional role for the conserved catalytic site in mammalian innate immune signaling.

## Introduction

Toll/Interleukin-1 receptor (TIR) domains, first characterized in animal Toll-like receptors (TLRs) and interleukin receptors, are immune modules conserved across the tree of life. TIR domains were originally described in animals as adaptors that recruit downstream signaling proteins via protein-protein interactions^1–3^. However, recent studies in bacteria and plants show that TIR domains are enzymes that produce immune signaling molecules using the molecule NAD^+^ as a substrate. In bacteria, proteins with TIR domains function within the Thoeris anti-phage system, where they serve to recognize phage infection^4,5^. Once infection is recognized, the bacterial TIR synthesizes an immune signal that contains ADP-ribose (ADPR), and this molecule activates a second protein within the Thoeris system that then causes growth arrest or premature cell death to abort phage infection^5^.

Signaling molecules produced by bacterial TIR proteins usually comprise a cyclical form of ADPR. These include 3ʹcADPR and 2ʹcADPR, molecules in which cyclization occurs via ribose-ribose bonds, as well as N7-cADPR and canonical cADPR, where cyclization involves ribose-adenine bonds^6–9^. A linear signaling molecule, in which ADPR is conjugated to histidine, was also described as produced by a certain Thoeris type in bacteria^10^. TIR-produced signals were shown to activate multiple kinds of bacterial immune proteins, including enzymes that deplete NAD^+^ to halt phage infection^5,11^, caspase-like proteases that cleave cellular and phage proteins^9^, and a variety of membrane-spanning proteins and ion channels that likely breach the membrane barrier when activated by the TIR-derived signal^8,10,12^.

In parallel with the discovery that bacterial TIRs generate signaling molecules, it was shown that plant TIR-containing immune receptors also catalyze the production of small-molecule signals^13–16^. Plant TIR domains commonly occur at the N-termini of intracellular immune proteins called nucleotide-binding leucine-rich repeat receptors (NLRs). These proteins sense pathogens via their C-terminal leucine-rich repeats, and then oligomerize, activating the TIR to produce the signaling molecule. The resulting signal triggers a complex containing the protein EDS1, which promotes downstream immunity that sometimes culminates in cell death^17–19^. Plant TIRs produce several different ADPR-containing signaling molecules, including 2ʹcADPR^6,7,20^, which is thought to be further processed in the plant cell before activating the EDS1-containing complex^21,22^. Some plant TIRs are thought to generate linear ADPR conjugates such as ADPr-ATP and di-ADPR^18,19^, as well as other products^15,18,19,23,24^.

TIR-mediated production of immune signaling molecules has not been reported in animals to date. Some animal TIR domains are known to have an NADase catalytic activity, but it is thought that their role is to deplete NAD^+^ rather than produce signaling molecules^13,25–27^. This is the case for the *C. elegans* protein TIR-1, whose NAD^+^ depletion activity is required for activation of the p38 immune pathway^26,28^. The only human TIR domain known to have a catalytic activity is encoded in the SARM1 protein, which acts in the nervous system to deplete NAD⁺ in response to axon injury^13,29^, and whose NADase activity was implicated in cytokine regulation in human monocytes^30^. In contrast, the TIR domains of TLRs and interleukin receptors are currently thought to function exclusively as protein-protein interaction modules^1–3^.

In the current study we set out to systematically examine the catalytic activities of TIR-domains across a broad range of eukaryotes, with a specific focus on animal TIRs. By experimentally testing >150 TIR domains, we show that TLR TIRs from multiple animals, including ones from the human and mouse TLR4, produce cADPR *in vitro*. We further demonstrate that the production of cADPR is a trait common to TIRs across the animal phylogenetic tree, suggesting that this molecule has a role in animal immune signaling. We further find that a point mutation in the active site of human TLR4 disrupts immune signaling in response to LPS stimulation and prevents Myddosome assembly. Finally, we show that a TIR from lancelet, a primordial chordate, produces the molecule 2ʹcADPR, while algal TIRs can produce 3ʹcADPR, suggesting conservation of these signaling molecules between bacteria, plants, and animals. Our study shows that enzymatic activity is a common, conserved trait of the TIR domain across the tree of life.

## Results

### Collection and phylogenetic analysis of eukaryotic TIRs

To comprehensively examine the functional and evolutionary diversity of TIR domains in eukaryotes, we retrieved ∼71,000 eukaryotic proteins annotated with a TIR domain from the UniProt database^31^ (Table S1). We first clustered the full-length proteins based on sequence similarity, and then extracted the TIR domain sequence from each of the proteins, yielding ∼16,000 TIR domains (Table S2). We then further clustered the TIR domains based on sequence homology, so that TIRs were clustered together if they shared 40% or more sequence identity (see Methods). After removing ‘singleton’ TIR sequences that did not cluster with any other TIR, this process generated about 1,500 clusters of eukaryotic TIRs, and one representative sequence from each cluster was taken for further analysis (Tables S3). We performed structure-guided phylogenetic analysis using the cluster representatives, yielding a phylogenetic tree spanning the diversity of eukaryotic TIRs (Figure 1A).

**Figure 1.**
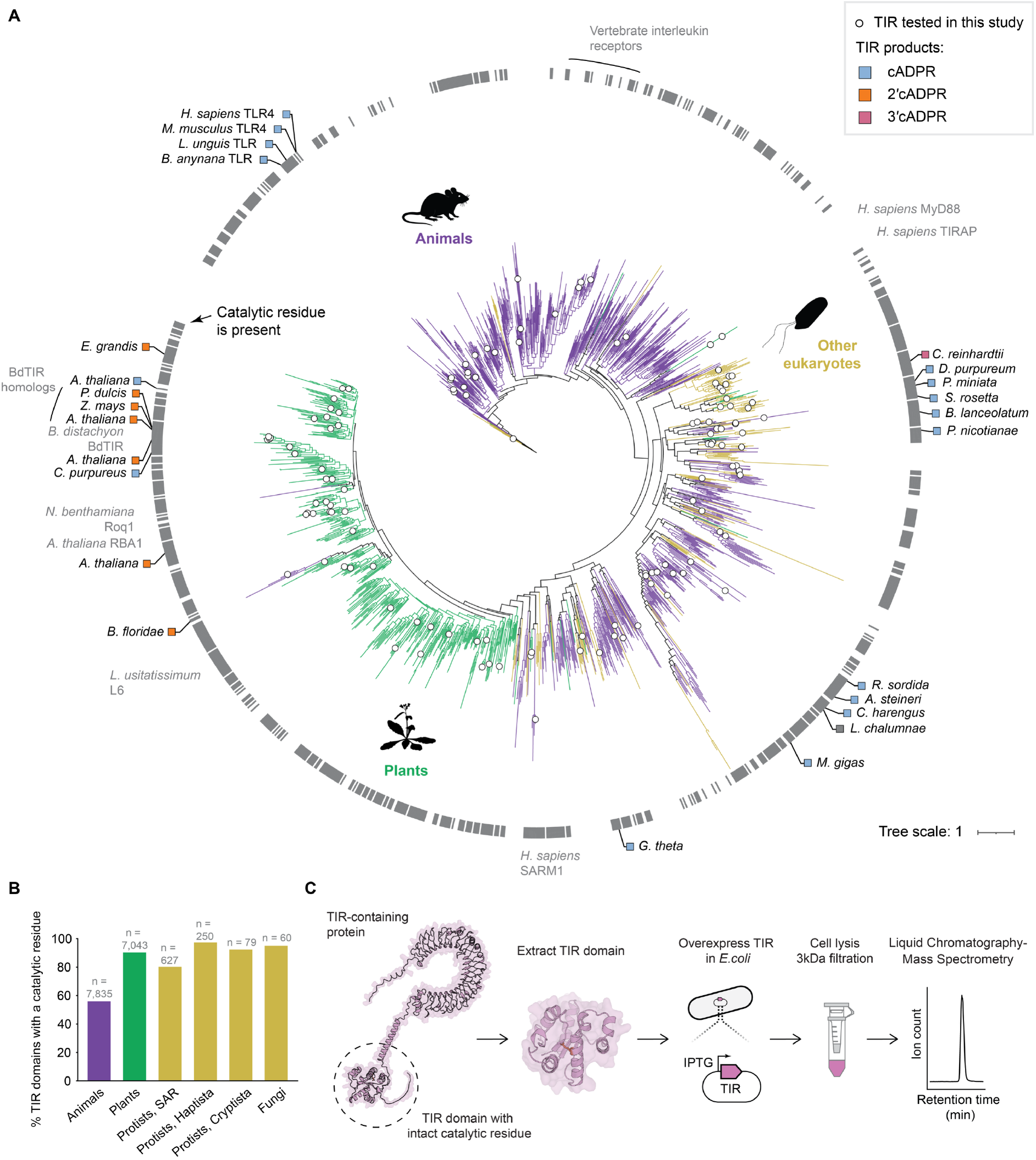
Diverse eukaryotic TIR-domain proteins contain conserved catalytic residues. **(A)** A phylogenetic tree of 1,522 representative eukaryotic TIR domains. Branch colors indicate the taxonomic group of the source organism. Grey tick marks in the outer ring indicate the presence of the conserved catalytic glutamic acid residue in the representative TIR. White circles on internal branches mark clusters from which one or more sequences were experimentally tested in this study. TIR domains that showed catalytic activity are indicated on the tree periphery in black, with the box color indicating their products. A TIR from *L. chalumnae* depleted NAD^+^ without generating a detectable cyclic ADPR product. Previously studied TIRs are indicated on the tree periphery in grey font. **(B)** Fraction of TIR-domain proteins that contain a glutamic acid residue in the catalytic position. Data are presented for eukaryotic clades in which >50 TIR-containing proteins exist in our dataset. **(C)** Schematic overview of the pipeline used in this study to identify eukaryotic TIR domains with catalytic activity.

TIR domain sequences were generally grouped on the tree based on their phyletic relationship. Almost all plant TIRs, for example, were grouped together in a single clade, suggesting that they have evolved from a small set of TIRs in the ancient ancestor of *Viridiplantae* (Figure 1A). Similarly, the TIRs of most animal TLRs and interleukin receptors clustered together in animal-specific clades (Figure 1A). These observations are in line with reports from earlier phylogenetic studies that were based on smaller subset of TIRs^32–35^.

A defining feature of catalytic TIRs that process NAD^+^ as a substrate is a conserved glutamic acid residue that protrudes from helix αC into the catalytic pocket (Figure S1). This glutamic acid residue is essential for the enzymatic activity of the TIR^13,14,29,36^, and numerous studies showed that a point mutation in this residue invariably abolishes the ability of TIRs to process NAD^+^ ^5,13,14,29,36^. We used structural alignments to determine the presence or absence of the catalytic residue in the ∼16,000 analyzed TIR domains (Table S2). As expected, the majority of TIR domains from animal TLRs and interleukin receptors lacked a catalytic site, in line with their known roles as protein-protein interaction scaffolding domains (Figure 1A). However, beyond these proteins, most TIRs in animals, plants, and other eukaryotes exhibited intact glutamic acid residue in the active site position, and even within the TLR clade, some subclades contained a conserved glutamate within the active site (Figure 1A, 1B). These results imply that many eukaryotic TIRs may be catalytically active.

### Experimental characterization of TIR domains

Given the presence of the catalytic site in many eukaryotic TIRs, we set out to experimentally test whether TIR domains from eukaryotic proteins possess an enzymatic activity. For this, we sampled the TIR tree for TIRs that have a conserved glutamate in the catalytic position of the TIR active site (Figure 1; Tables S2, S3). The sampled TIRs included 61 TIRs from animals, 29 TIRs from non-animal unicellular eukaryotes, and 13 TIRs from green algae and plants that were not localized to the main clade of plant TIRs. We also included 51 additional TIRs from the main clade of plants and green algae that were not previously tested for catalytic activities. Overall, this set included 154 TIRs from 96 evolutionarily diverse species (Table S4).

It was previously shown that, when overexpressed in *Escherichia coli*, some immune TIR domains can generate their cognate signaling molecules^14,36^. Notably, overexpression experiments of plant TIRs in *E. coli* were the basis for the original discovery that plant TIRs produce immune signals^14^. To test if the TIR domains we collected possess an enzymatic activity, we overexpressed each of the 154 TIRs in *E. coli*, and then analyzed the cell lysates using a liquid chromatography-mass spectrometry (LC-MS) protocol tailored to detect three cyclic ADPR isomers previously reported to be generated by TIR domains^13–15^ (Figure 1C; Figure S2). When the TIR domain was encoded on a relatively short protein without additional domains, we expressed the full-length protein (25 TIRs, ∼16% of all cases), whereas for the rest of the proteins we expressed the TIR domain only (Table S4).

We verified, using a set of control TIR domains known to produce signaling molecules, that our process enables the identification of the cognate molecules produced by TIR domains. For this, we tested the verified control plant TIR proteins BdTIR, L6, Roq1, and RBA1, and the bacterial protein AbTIR, all of which are known to produce 2ʹcADPR^6,7,14^. We also similarly tested the bacterial proteins HopAM1 and ThsBʹ, known to produce 3ʹcADPR^7,37,38^. Our experiments verified that the positive control TIRs produced the expected molecules in the cell lysate, validating our protocol for LC-MS detection of TIR-derived signaling molecules using heterologous expression in bacteria (Figure S3).

Our data show that a non-negligible fraction of the tested TIRs produced one of the three tested cyclic ADPR isomers (Figure 1A; Table S4). Of the 61 animal TIRs tested, 12 showed catalytic activity, with 4 TIRs from non-animal unicellular eukaryotes showing activity as well. Nine plant and green algal TIRs also produced molecules that were detectable in the lysates (Table S4). As expected^7,20,24^, the majority of active plant TIRs produced 2ʹcADPR (Figure S4), but surprisingly, a TIR domain from the green alga *Chlamydomonas reinhardtii* produced substantial amounts of 3ʹcADPR (Figures S5, S6, S7). Most of the active animal TIRs, however, produced the canonical cADPR molecule, again with some exceptions that are described in more detail below. We were unable to detect His-ADPR and pRib-AMP in TIR-expressing cells, suggesting these compounds are not produced by the TIR domains we studied (Figures S8, S9).

### Human TLR4 TIR domain produces cADPR when expressed in bacteria

Surprisingly, one of the TIR domains that generated cADPR when expressed in bacteria was the TIR from human TLR4 (Figure 2). This protein has a well-documented, central role in human antibacterial immunity, where it activates an innate immune response once it senses bacteria-derived lipopolysaccharides (LPS)^3,39,40^. However, to date, there have been no reports indicating that the TIR domain of TLR4, or any other TLR, has an enzymatic capacity.

**Figure 2.**
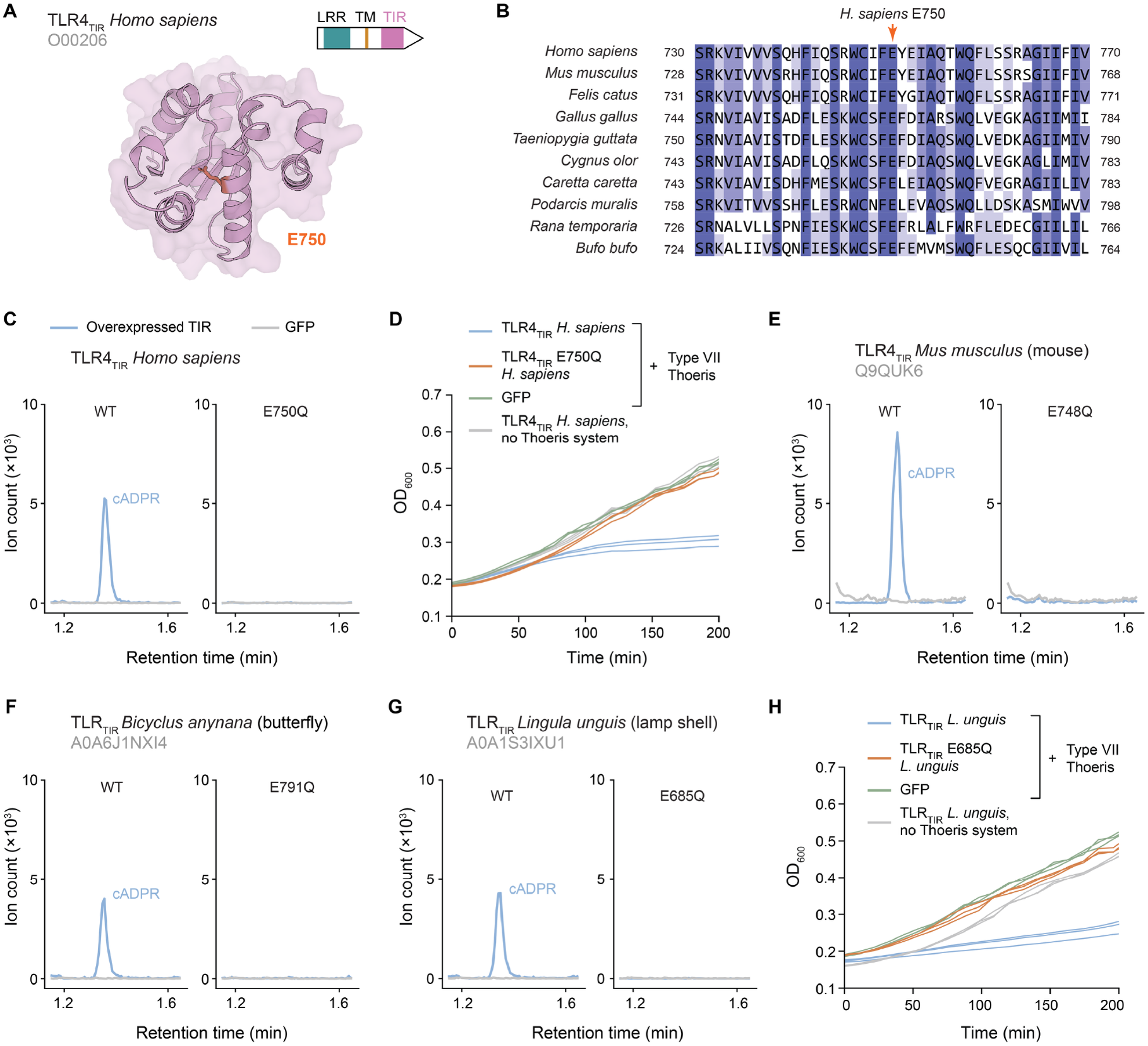
The TIR domain of TLR4 produces cADPR. **(A)** AlphaFold3^54^-predicted structure of the *Homo sapiens* TLR4 TIR domain, with the catalytic glutamic acid residue (E750) highlighted in orange. The domain architecture of the full-length TLR4 is schematically shown above. **(B)** The catalytic residue is conserved among orthologs of TLR4. Multiple sequence alignment of TLR4 TIR domain orthologs, showing a window of residues corresponding to amino acids 730-770 in the human TLR4 protein. Conserved residues are highlighted in purple. Source species are indicated on the left. **(C)** TIR-dependent production of cADPR. Extracted ion chromatograms (m/z = 540.053) from lysates of *E. coli* cells expressing the wild-type TLR4 TIR domain, or a catalytic mutant (E750Q), for 1 hour. Data are representative of three replicates. **(D)** Human TLR4 TIR is toxic when co-expressed with type VII Thoeris. Growth curves of *E. coli* cells co-expressing the wild-type TLR4 TIR domain, a catalytic mutant (E750Q) of the TLR4 TIR, or GFP, together with a type VII Thoeris system cloned from the metagenomic *Alteromonadales* scaffold^8^. Three repeats are shown as individual curves. **(E-G)** Extracted ion chromatograms (m/z = 540.053) from lysates of *E. coli* cells expressing wild-type or catalytic mutant TIR domains from **(E)** *Mus musculus* TLR4, TIR expression was induced for 4 hours; **(F)** *Bicyclus anynana* (butterfly), TIR expression was induced for 1 hour; and **(G)** *Lingula unguis* (brachiopod), TIR expression was induced for 1 hour. **(H)** Growth curves of *E. coli* cells co-expressing wild-type or catalytic mutant (E685Q) *L. unguis* TIR domain or GFP together with the type VII Thoeris system or RFP. Three repeats are shown as individual curves. The GFP control LC-MS data (1 hour induction) shown in panels C, F, and G are also shown in Figures 3B, S3, S4, S5, S11, S13, and S15; GFP control data (4 hours induction) presented in panel E are also shown in Figure 3B. All three repeats of GFP control for both 1-hour and 4-hour induction are shown in Figure S2B and C. Data for control GFP expressed together with type VII Thoeris presented in panel D, and H are also shown in Figure 3C.

We found that the indicative glutamic acid residue in the catalytic position of the TLR4 TIR is invariably conserved across TLR4 homologs extending across all vertebrates (Figure 2A, 2B). While cADPR was detected in bacteria expressing the TLR4 TIR, a mutated form of TLR4 TIR, in which a point mutation was introduced to substitute the glutamic acid residue with glutamine (E750Q), did not produce cADPR when expressed in bacteria, providing further support that cADPR production depended on the catalytic activity of the TIR (Figure 2C).

A recent study revealed a bacterial Thoeris system in which the TIR domain produces cADPR in response to infection^8^. In this system, called type VII Thoeris, the cADPR molecule activates a membrane-spanning protein that impairs membrane integrity and halts bacterial growth^8^. To further substantiate that the molecule produced by the human TLR4 TIR is indeed the canonical cADPR, we expressed that TIR in an *E. coli* strain that also expressed type VII Thoeris. A marked growth inhibition was observed in cells co-expressing the TLR4 TIR and the Thoeris system, but not in cells expressing the mutated TIR (E750Q) or the WT TIR in the absence of Thoeris (Figure 2D). These results suggest that the TIR-produced cADPR activated the toxic effector of type VII Thoeris. As the effector of type VII Thoeris is specifically activated only in response to canonical cADPR but not by any other known TIR-generated cyclic ADPR isomer^8^, these results verify the identity of the molecule produced by the TLR4 TIR.

Beyond the TIR from human TLR4, three additional TLR TIRs in our set produced cADPR when expressed in bacteria. One of these was the TIR from the mouse TLR4 (Figure 2E), which displays 67% sequence identity when compared to the human TLR4 protein sequence. The conservation of cADPR production between human and mouse TLR4 TIRs lends further support to the hypothesis that catalytic activity is a conserved feature in TLR4 biology. TIRs from two invertebrate TLRs, the butterfly *Bicyclus anynana* and the brachiopod *Lingula unguis*, also produced cADPR (Figure 2F, 2G). Our data show that TIRs from invertebrate TLRs are also capable of activating Thoeris-mediated toxicity, in a manner dependent on the predicted catalytic glutamate (Figure 2H). Together, these results indicate that cADPR-producing enzymatic activity is conserved among TLR TIR domains across both vertebrates and invertebrates.

### TIR domains from evolutionarily diverse species produce cADPR

Beyond the unexpected catalytic activity observed in the TIR domain of TLR4 proteins, the majority of animal TIR domains exhibiting cADPR production were localized on the phylogenetic tree outside of the Toll-like and interleukin receptor clades (Figure 1A). Among those were proteins with no other domain except for the TIR domain (reminiscent of the plant immune protein BdTIR), or TIRs found in proteins that encode other immune-associated domains, consistent with a role in immune signaling (Figure 3A). This distribution suggested that immune-related enzymatic activity among animal TIR domains may be substantially more widespread than manifested by TLRs alone.

**Figure 3.**
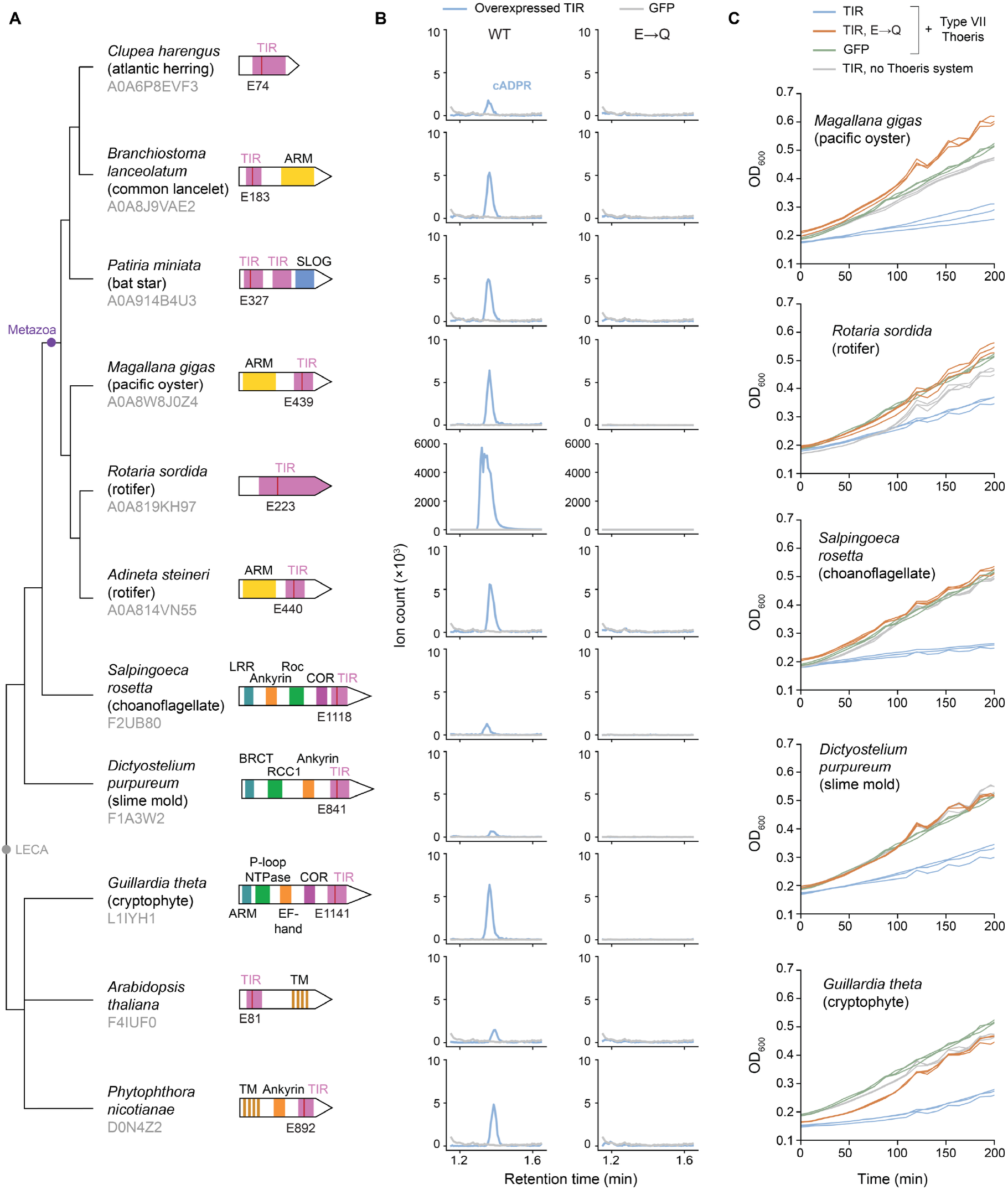
TIR domains from evolutionarily diverse eukaryotes produce cADPR. **(A)** Cladogram of eukaryotic species whose proteins are shown in this figure, with the last eukaryotic common ancestor (LECA) and Metazoa clades indicated. Domain architectures of the full-length TIR-containing proteins are shown to the right. **(B)** Extracted ion chromatograms (m/z = 540.053) from lysates of *E. coli* cells expressing wild-type or catalytic mutant variants of the TIR domains indicated in panel A. Induction times for each TIR domain (one or four hours) are listed in Table S4. Data are representatives of three replicates. **(C)** Growth curves of *E. coli* cells co-expressing the indicated eukaryotic TIR domains or GFP with the type VII Thoeris system or an empty vector. Three repeats are shown as individual curves. Abbreviations: ARM, armadillo; BRCT, BRCA1 C-terminal; COR, C-terminal of Roc; LRR, leucine-rich repeat; NTPase, nucleoside-triphosphatase; RCC1, regulator of chromosome condensation 1; Roc, Ras of complex; SLOG, SMF/DprA-LOG domain; TM, transmembrane. The GFP control LC-MS data (1 hour induction) shown in panel B are also shown in Figures 2C, 2F, 2G, S3, S4, S5, S11, S13, and S15; GFP control data (4 hours induction) are shared with Figure 2E. All three repeats of GFP control for both 1-hour and 4-hour induction are shown in Figure S2B and C. The control data for GFP expressed together with type VII Thoeris in panel C are also shown in Figures 2D and 2H.

The TIRs found to produce cADPR spanned a wide range across distantly related animal lineages, including both protostomes (mollusk, rotifers) and deuterostomes (fish, echinoderm, lancelet), suggesting that this catalytic activity is broadly conserved in Metazoa (Figure 3A). cADPR-producing TIR domains were also detected outside Metazoa, in phylogenetically diverse unicellular eukaryotes including the choanoflagellate *Salpingoeca rosetta*, the amoebozoan *Dictyostelium purpureum*, the cryptophyte *Guillardia theta*, and the oomycete *Phytophthora nicotianae* (Figure 3A, 3B, 3C). In all cases, TIR activity was dependent on the existence of the catalytic glutamate, and a point mutation in that residue abolished cADPR production (Figure 3B). AlphaFold3 placed cADPR within the catalytic pocket in a consistent orientation in all cases (Figure S10). As observed for the TLR4 TIR, non-TLR TIRs activated type VII Thoeris-mediated growth arrest when expressed in bacteria (Figure 3C).

Some catalytic TIR domains are known to deplete NAD^+^ rather than produce signaling molecules^25,29,41^. NAD^+^-depleting TIRs usually convert NAD^+^ to linear ADPR, although in some cases, such as SARM1 from human and *Drosophila*, NAD^+^ depletion results in the production of a mixture of ADPR and cADPR as major products, with additional minor products^29,42^ (Figure S11). Although we cannot rule out that the role of some of the catalytic TIRs described here is to deplete NAD^+^ rather than produce cADPR as a signaling molecule, we note that in most cases, NAD^+^ was not depleted in cells expressing the TIRs we studied, even after overnight overexpression of the TIR (Figure S12). Notable exceptions included a TIR domain from the fish *Latimeria chalumnae* that depleted cellular NAD⁺ without producing any of the cADPR isomers we measured (Table S4), and TIR domains from the cryptophyte *Guillardia theta*, the rotifer *Rotaria sordida*, and the moss *Ceratodon purpureus*, which produces cADPR (Figure 3B; Figure S13) and depleted cellular NAD^+^ after overnight expression (Figure S12). The *C. purpureus* TIR also produced detectable minor levels of 2ʹcADPR and 3ʹcADPR alongside the primary cADPR (Figure 13), and biochemical assays^9^ showed that it can also produce the unstable molecule N7-cADPR (Figure S14). Another TIR whose overnight expression led to NAD^+^ depletion was one from the lancelet *Branchiostoma floridae* (an early-diverging chordate). This TIR produced detectible amounts of 2ʹcADPR but none of the other cADPR isomers we measured (Figure S15).

### The catalytic residue of TLR4 is required for LPS-induced signaling in human macrophages

Our finding that the TIR domain of human TLR4 possesses catalytic activity prompted us to investigate the contribution of its catalytic glutamate to TLR4 signaling in human cells. Using CRISPR/Cas-mediated genome editing and a homology-directed repair template, we introduced the E750Q substitution into the endogenous *TLR4* locus in CEBRA cells, a human cellular model in which B cells can be transdifferentiated into macrophages by forced expression of C/EBPα in the presence of IL-3 and M-CSF^43^. We recovered several independent clones in which sequencing detected only the edited allele (Figure 4A). Following transdifferentiation into macrophages, TLR4 protein was readily detectable in these clones, although its abundance was moderately lower than in wild-type cells (Figure 4B). Stimulation with LPS induced robust production of IL-6 and CXCL10 (IP-10) in wild-type macrophages, whereas production of both cytokines was not detected in E750Q TLR4 macrophages (Figure 4C). In contrast, stimulation with the TLR7/8 agonist R848 elicited comparable cytokine production in wild-type and E750Q cells, indicating that the mutant cells retained responsiveness to an independent innate immune stimulus (Figure 4C).

**Figure 4.**
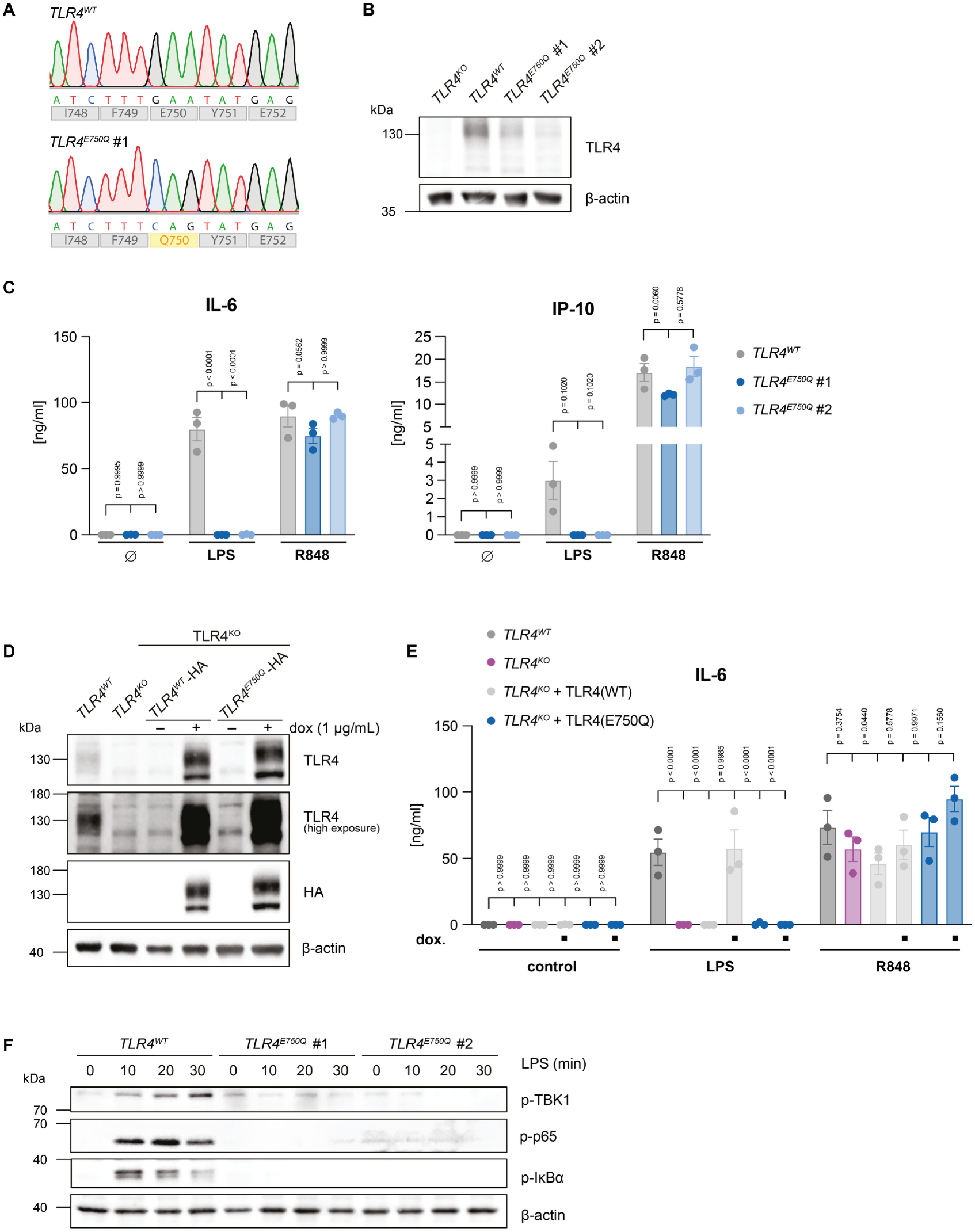
The catalytic residue of TLR4 is required for LPS-induced signal transduction in human macrophages. **(A)** Sanger sequencing chromatograms of the endogenous *TLR4* locus in wild-type (WT) CEBRA cells and one exemplary independently generated *TLR4* E750Q knock-in clone. **(B)** Western blot analysis of TLR4 expression in WT cells and the two *TLR4* E750Q knock-in clones. β-actin served as a loading control. One representative experiment of two independent experiments is shown. **(C)** WT cells and two independent *TLR4* E750Q knock-in clones were stimulated with LPS or R848. IL-6 and IP-10 concentrations in the culture supernatants were measured by ELISA 16 hours after stimulation. Data are presented as mean ± SEM of three independent experiments. Statistical significance was determined by two-way ANOVA followed by Dunnett’s multiple-comparisons test. **(D)** Western blot analysis of CEBRA cells of the indicated genotypes reconstituted with doxycycline-inducible WT or E750Q TLR4. TLR4 expression was analyzed in the absence or presence of doxycycline. One representative experiment of three independent experiments is shown. **(E)** Cells reconstituted with WT or E750Q TLR4 were treated with doxycycline (1 µg/mL) for 6 hours, followed by stimulation with LPS or R848, and IL-6 concentrations in the culture supernatants were measured by ELISA 16 hours after stimulation. Data are presented as mean ± SEM of three independent experiments. Statistical significance was determined by two-way ANOVA followed by Dunnett’s multiple-comparisons test. **(F)** Western blot analysis of NF-κB pathway activation in WT cells and two independent *TLR4* E750Q knock-in clones following stimulation with LPS for the indicated times. Phosphorylation of TBK-1, p65, and IκBα was assessed, while β-actin served as a loading control. One representative experiment of three independent experiments is shown.

Although the moderate reduction in TLR4 abundance seemed unlikely to account for the complete loss of cytokine production, we directly addressed this possibility by reconstituting TLR4 knockout cells with either HA-tagged wild-type TLR4 or the E750Q variant using a doxycycline-inducible expression system. Western blot analysis confirmed that both proteins were expressed at levels substantially higher than endogenous TLR4 (Figure 4D). Reconstitution with wild-type TLR4 restored LPS-induced IL-6 production, whereas cells expressing the E750Q variant remained completely unresponsive (Figure 4E). Responses to R848 were preserved in both populations. Thus, even expression substantially above endogenous levels failed to restore signaling by E750Q TLR4, arguing against reduced receptor abundance as the explanation for the mutant phenotype.

To determine where the E750Q substitution disrupts transcriptional response to LPS, we examined induction of inflammatory gene expression. LPS induced *IL6* and *TNF* transcripts in wild-type macrophages, whereas this response was completely abrogated in macrophages carrying the endogenous E750Q substitution (Figure S16A). We confirmed this defect in the reconstitution system: LPS induced *IL6* and *TNF* transcripts in TLR4 knockout cells reconstituted with wild-type TLR4, but not in TLR4 knockout cells or cells reconstituted with the E750Q variant (Figure S16B). To distinguish a population-wide loss of responsiveness from a reduction in the proportion of responding cells, we generated a transcriptional reporter by inserting a T2A-mScarlet cassette at the endogenous *IL1B* locus, one of the most strongly inducible NF-κB target genes in CEBRA cells. Following LPS stimulation, both wild-type cells and TLR4 knockout cells reconstituted with wild-type TLR4 showed a population-wide shift to mScarlet positivity. In contrast, TLR4 knockout cells and cells reconstituted with E750Q TLR4 remained unresponsive (Figure S16C and D). R848 activated the reporter comparably in all four cell populations, demonstrating that their capacity to induce inflammatory gene expression remained intact (Figure S16C and D). These results place the E750Q-associated defect at or upstream of inflammatory gene induction.

We next examined activation of the canonical NF-κB pathway by monitoring phosphorylation of its inhibitor IκBα, the NF-κB subunit p65, and the kinase TBK-1. In wild-type macrophages, LPS-induced IκBα phosphorylation was detectable as early as 10 minutes, whereas p65 phosphorylation showed a slightly delayed onset and peaked at 20–30 minutes. Phosphorylation of both proteins was completely abrogated in macrophages carrying the endogenous E750Q substitution (Figure 4F). We confirmed this phenotype in the reconstitution system: LPS-induced phosphorylation of IκBα and p65 was observed in wild-type cells and in TLR4 knockout cells reconstituted with wild-type TLR4, but not in TLR4 knockout cells or cells reconstituted with E750Q TLR4 (Figure S16E). Together, these findings show that the E750Q substitution disrupts signaling upstream of canonical NF-κB activation, preventing subsequent inflammatory gene induction and cytokine production.

### The catalytic residue of TLR4 is required for Myddosome assembly

We next asked whether the requirement for E750 persists when the intracellular signaling module of TLR4 is activated independently of LPS recognition. To address this, we used a chimeric receptor in which the extracellular domain of CD16 was fused to the intracellular signaling region of TLR4^44^ (Figure 5A). Doxycycline-inducible expression of the wild-type and E750Q chimeras yielded comparable protein levels (Figure 5B). Induction of the wild-type chimera elicited robust IL-6 and IP-10 production (Figure 5C) and activation of the IL1B^mScarlet^ reporter (Figure S17A and B). In contrast, at low expression levels, the E750Q chimera elicited little or no IL-6 production or reporter activation, and IP-10 production remained undetectable. At higher doxycycline concentrations, the E750Q chimera induced partial IL-6 production and IL1B^mScarlet^ reporter expression, reaching approximately one third of the responses observed with the wild-type chimera (Figure 5C; Figure S17A and B). This partial recovery suggests that high expression of the chimera can partially bypass the requirement for E750 for these readouts. Nevertheless, signaling remained substantially impaired, supporting a role for E750 within the intracellular signaling module of TLR4 independently of LPS recognition.

**Figure 5.**
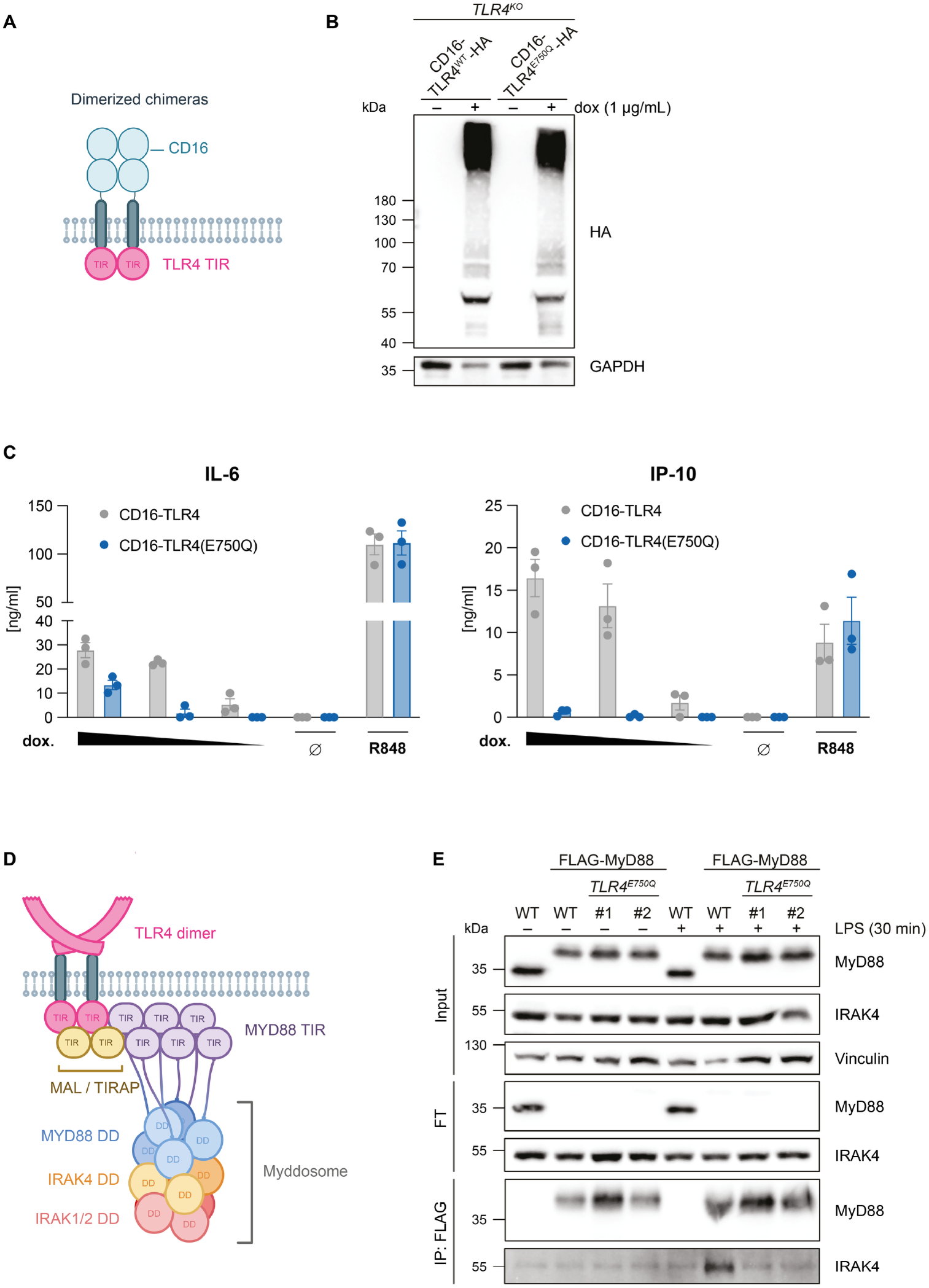
The catalytic residue of TLR4 is required for proximal TLR4 signal transduction and Myddosome assembly. **(A)** Schematic representation of the CD16–TLR4-TIR chimeric receptor used to induce dimerization of the intracellular TLR4 signaling domain independently of LPS recognition. **(B)** Western blot analysis of *TLR4* knockout CEBRA cells expressing doxycycline-inducible CD16–TLR4-TIR constructs containing either the WT or E750Q TLR4 TIR domain. Expression of the constructs was detected using the HA tag, and GAPDH served as a loading control. One representative experiment of three independent experiments is shown. **(C)** CEBRA cells expressing doxycycline-inducible WT or E750Q CD16–TLR4-TIR constructs were treated with 37, 12, or 4 ng doxycycline, left untreated, or stimulated with R848 as a control. IL-6 and IP-10 were measured by ELISA 24 hours after induction. Data are presented as mean ± SEM of three independent experiments. **(D)** Schematic representation of TLR4-induced Myddosome assembly, showing recruitment of MyD88 and IRAK4 downstream of activated TLR4. **(E)** Analysis of LPS-induced Myddosome assembly by FLAG immunoprecipitation. WT CEBRA cells or cells carrying an endogenous N-terminal FLAG tag on MyD88 in either a *TLR4* WT or E750Q knock-in background were left untreated or stimulated with LPS. Input, flow-through (FT), and FLAG immunoprecipitation fractions were analyzed by Western blot for the indicated proteins. Two independent *TLR4* E750Q FLAG knock-in clones were used for the experiment. One representative experiment of three independent experiments is shown.

We therefore examined whether the E750Q substitution disrupts assembly of the Myddosome, the proximal signaling complex containing MyD88 and IRAK family kinases (Figure 5D). To assess this in a setting that preserves endogenous expression of both receptor and adaptor, we introduced an N-terminal FLAG tag at the endogenous *MYD88* locus in wild-type cells and one TLR4 E750Q knock-in clone. Following LPS stimulation for 30 minutes, we immunoprecipitated MyD88 and assessed its association with the Myddosome component IRAK4. Comparable amounts of MyD88 were recovered from wild-type and E750Q cells. However, whereas LPS induced robust association of IRAK4 with MyD88 in wild-type cells, IRAK4 recruitment was undetectable in both E750Q clones (Figure 5E). Thus, the E750Q substitution disrupts TLR4 signaling at or upstream of Myddosome assembly, placing the requirement for the conserved catalytic glutamate at an early step in intracellular signal transduction.

Together, these results establish that the conserved catalytic glutamate of the TLR4 TIR domain is essential for LPS-induced proximal signal transduction in human macrophages. Combined with our biochemical findings, these data support a functional role for TIR enzymatic activity in human TLR4 signaling.

## Discussion

Our findings reveal that the catalytic activity of TIR domains in eukaryotes is far more widespread than previously appreciated. While this was expected for plant TIRs due to their recent characterization as immune signal producers^14,19^, our findings that animal TLR TIRs can produce cADPR molecules are surprising given the general perception of these domains as protein–protein interaction modules^1,3^. Our observation that the indicative catalytic glutamate essential for TIR NADase activity^14,29,36^ is conserved among eukaryotic TIRs across diverse clades, suggests that the enzymatic activity is the ancestral function of the TIR domain.

The catalytic activity we measured for the TLR4 TIR is especially surprising given that the TLR4 pathway has been extensively studied in human immunology^3,39^. Importantly, our experiments in human macrophages establish that the conserved catalytic residue of TLR4 is functionally required for LPS-induced signaling, as substitution of E750 with glutamine abolished LPS-induced cytokine production and NF-κB activation. The signaling defect persisted when WT and E750Q TLR4 were expressed at comparable levels and when the intracellular TLR4 signaling module was activated independently of LPS recognition.

When TLR4 is activated by LPS, the adapter protein MAL (also called TIRAP) is recruited to the TLR4 TIRs^3^. MAL then recruits MyD88 to nucleate Myddosome assembly^45,46^. E750Q TLR4 failed to support LPS-induced MyD88-IRAK4 association, placing the defect at or upstream of Myddosome assembly, the earliest stage of TLR4 signaling that we could directly examine in our cellular system. The proximal nature of the signaling defect provides a clue to the possible function of the TLR4 catalytic site. Based on the structure of MAL TIR-domain filaments, Ve et al. proposed that activated TLR4 TIR domains associate through a head-to-tail intrastrand interface, generating a platform that recruits MAL/TIRAP through a distinct interstrand interface^46^. The region surrounding E750 lies at the modeled TLR4-MAL interstrand interface. Although this model does not establish E750 itself as a direct MAL-contact residue, its location, together with the loss of Myddosome assembly in the E750Q mutant, raises the possibility that TLR4 NAD^+^ processing is functionally coupled to MAL recruitment or organization of the receptor-proximal TIR complex. We could not directly examine MAL recruitment in our cellular system, and this hypothesis therefore remains to be tested in a future study.

Notably, IP-10 production was also essentially abolished in E750Q cells. Because induction of this response involves the TRAM-TRIF branch of TLR4 signaling^47^, this observation raises the possibility that disruption of the catalytic site affects the two major TLR4 adaptor pathways. Direct analysis of TRAM/TRIF signaling will be required to establish this point; nevertheless, impairment of both MyD88-dependent responses and IP-10 production is consistent with a defect occurring very early in the signaling cascade, potentially at the level of TIR-dependent adaptor recruitment or organization.

This connection is particularly intriguing considering the mechanism of the TLR4-selective inhibitor TAK-242 (resatorvid). TAK-242 covalently binds C747^48^, a residue residing near the E750 NAD^+^ processing region, and inhibits TLR4 signaling. Previous work further showed that TAK-242 disrupts association of TLR4 with both TIRAP/MAL and TRAM^49^ placing its inhibitory effect at a receptor-proximal stage similar to that implicated by our E750Q experiments. Although these observations have been interpreted in terms of disruption of TLR4-adaptor interactions, the position of C747 near the site implicated in NAD^+^ processing raises an alternative, but not mutually exclusive, possibility that covalent modification by TAK-242 perturbs catalytic-site function. Although the structure of TAK-242 in complex with TLR4 TIR was not determined, AlphaFold3 places TAK-242 with high confidence in the predicted catalytic pocket of the TLR4 TIR in a manner that would block access to the NAD^+^ substrate (Figure S18). Determining whether TAK-242 directly inhibits the NAD^+^-processing activity of the TLR4 TIR domain will be important for establishing whether pharmacological inhibition by TAK-242 and mutation of the catalytic residue converge on a common receptor-proximal mechanism.

In addition to TLR4, other human TLRs and interleukin receptors have a conserved glutamate at the catalytic position of the TIR active site^13,50^. Previous studies, which attempted to test NADase activity of multiple TIR domains, including those of IL-1R9, IL-1R10, and TLR2, did not detect such an activity *in vitro* despite the presence of the indicative glutamate residue in these TIRs^13,50^. Analysis of the crystal structures of these three TIRs indicated that the glutamate residue is oriented in a way that faces out of the catalytic pocket, explaining the lack of NAD^+^ cleavage^50^. Consistent with these reports, the TIR domain of TLR2 was included in our set of tested TIRs and did not produce any cADPR isomer under our experimental conditions (Table S4). However, we note that it is possible that during TIR activation, conformational changes might bring the catalytic residue into a position favorable for NAD^+^ catalysis.

TIR-domain proteins have been identified in diverse unicellular eukaryotes through bioinformatic analyses^51^, and in *Dictyostelium discoideum* the TIR-domain protein TirA was shown to be required for immune-like sentinel cell function and for growth on live bacteria^52^. TIR-domain proteins were also recently implicated in antiviral defense in *Gephyrocapsa huxleyi*^53^. Our demonstration that TIR domains from representatives of Amoebozoa, Choanoflagellata, Cryptophyta, Oomycota, and Chlorophyta are catalytically active NADases opens a new direction for discovering immune mechanisms across the eukaryotic tree of life. Investigation of the physiological role of these TIRs could reveal cyclic nucleotide-based antiviral pathways in unicellular eukaryotes that may resemble those operating in bacteria, plants, and possibly animals.

Taken together, our results suggest that the catalytic activity of TIR domains plays a role in animal innate immunity. A few earlier observations hinted at this possibility: an NAD⁺-processing TIR-domain protein in *C. elegans* was implicated in antimicrobial defense^26^, and a TIR-STING protein in the mollusk *C. gigas* was suggested to deplete NAD⁺ in response to cGAS-like signaling^25^. As our results are mainly derived from overexpression of TIR domains in bacteria, they are, at this stage, speculative. However, we note that similar observations in bacteria and in plants initiated a revolution in understanding plant and bacterial immunity^5,13,14^. In this context, it is tempting to speculate that future studies exploring TIR enzymatic activities will drive another transformation in our understanding of animal immune signaling.

## Limitations of the study

While our study allowed screening of a large number of TIR domains for the production of cADPR, 2ʹcADPR and 3ʹcADPR, it has several major limitations. First, several molecules associated with plant TIRs signaling, including pRib-ADP, di-ADPR, and ADPr-ATP^18,19^, were not measured in our study because chemical standards were not available at the time our experiments were performed. We also did not quantify ADPR, the linear product of NAD^+^ hydrolysis, as this metabolite is naturally present in *E. coli* cells, masking ADPR signals in cell lysates. It is therefore possible that some of the TIRs for which we did not report catalytic activity are in fact enzymes that produce molecules not tested in our study. Another major limitation is that we measured the activity of TIR domains in isolation, outside their native protein context. While the *E. coli* overexpression system is well-established for detecting enzymatic activities of some TIRs^14,36^, it does not fully recapitulate physiological activation mechanisms, including oligomerization-dependent activation and auto-inhibition by flanking domains. The activities we report therefore reflect the catalytic capability of each TIR domain rather than its regulated activity in the source organism, and the conditions under which these domains are activated in their native context remain to be determined. Relatedly, enzymatic activity was determined for most domains from detection of the produced molecules and NAD^+^ depletion in cells rather than by direct measurement with purified protein, and we therefore do not report kinetic parameters for these domains. The cellular processes that respond to TIR-derived cADPR in animals, and to 3ʹcADPR in green algae, remain to be identified, as do the proteins that receive these signals. Finally, we note that due to the extreme sequence and structural divergence among eukaryotic TIR domains, bootstrap values are low for many of the internal nodes of the TIR phylogenetic tree (Figure 1A). We therefore used the tree as a framework for identifying major functional clades and guiding experimental selection, rather than as a precise reconstruction of TIR evolutionary history.

Although we established that the human TLR4 TIR domain produces cADPR when expressed in bacterial cells, we have not demonstrated TLR4-dependent NAD^+^ processing or product formation in human myeloid cells. Thus, while the biochemical activity of TLR4 and the cellular requirement for E750 connect the catalytic site to receptor function, they do not establish that loss of NAD^+^ processing causes the E750Q signaling phenotype. Direct measurement of TLR4-dependent NAD^+^ turnover and its products following receptor activation will be required to establish this link. Endogenous TLR4 abundance was moderately reduced in E750Q cells, raising the possibility that altered receptor stability contributes to impaired signaling. However, E750Q remained strongly defective when WT and mutant TLR4 were expressed at comparable levels in TLR4-deficient cells, as well as in the CD16–TLR4-TIR system, arguing against reduced receptor abundance as the primary explanation for the near-complete signaling defect. Nevertheless, an effect of E750Q on receptor stability or turnover cannot be fully excluded. More fundamentally, E750Q could perturb TIR-domain structure or TIR:TIR interactions independently of its effect on catalysis. This possibility is particularly difficult to exclude because the TLR4 TIR domain has thus far escaped high-resolution structural characterization, and TLR4 signaling depends on cooperative interactions between receptor and adaptor TIR domains. Although E750Q abolishes cADPR production by the isolated TLR4 TIR domain and disrupts signaling at or upstream of Myddosome assembly, these observations cannot formally distinguish loss of catalytic activity from a direct structural effect of the substitution. Biochemical and structural comparison of WT and E750Q TLR4, including their folding, self-association, NAD^+^ binding and processing, and interactions with TIR-containing adaptors, will therefore be important to establish whether catalysis itself is required for TLR4 signalosome formation.

## Resource availability

### Lead contact

Further information and requests for resources and reagents should be directed to the lead contacts, Rotem Sorek and Veit Hornung.

### Materials availability

This study did not generate new unique reagents.

### Data and code availability

Data that support the findings of this study are available in the article and Tables S1-S8. Any additional information required to reanalyze the data reported in this paper is available from the lead contact upon reasonable request.

## Acknowledgements

We thank members of the Sorek and Hornung labs for constructive discussions during this study. R.S. was supported, in part, by the European Research Council (grant ERC-AdG GA 101018520), the Israel Science Foundation (MAPATS grant 2720/22), the Deutsche Forschungsgemeinschaft (SPP 2330, grant 464312965), the Minerva Foundation with funding from the Federal German Ministry for Education and Research, a Max Planck-Humboldt Research Award, a research grant from Magnus Konow in honor of his mother Olga Konow Rappaport, and the Center for Immunotherapy at the Weizmann Institute of Science. V.H. was supported by grants from the Deutsche Forschungsgemeinschaft (DFG, German Research Foundation) CRC 1403/A03 (Project-ID 414786233), the European Research Council (ERC-2020-ADG–101018672 ENGINES), and the Cluster for Nucleic Acid Sciences and Technologies – NUCLEATE (DFG, Project-ID 533767322 – EXC 3113/1). I.O. was supported by the Ministry of Absorption New Immigrant program. E.Y. was supported by the Clore Scholars Program and, in part, by the Israeli Council for Higher Education (CHE) via the Weizmann Data Science Research Center. O.R. was supported by the Weizmann Postdoctoral Excellence Fellowship.

## Author contribution

B.H., S.B.C., V.H., and R.S. conceptualized and led the study. B.H. built and executed the computational pipeline, selected candidates for the screen, conducted *E. coli* toxicity experiments, and collected samples for mass spectrometry experiments with the help of J.G. I.O. assisted with planning LC-MS and *E. coli* toxicity experiments, and purified *C. reinhardtii* TIR domain. S.B.C. designed and conducted the experiments involving human cells and engineered the cell lines. S.J.S. and A.-L. B assisted with the experiments, and S.J.S. additionally conducted the imaging studies. A.H.F. conducted mass spectrometry experiments, and the data were processed together with B.H with the help of O.R. E.Y. identified novel Thoeris systems and assisted with building the computational pipeline. A.L. integrated the Type VII Thoeris system in the *E. coli* genome. G.A. assisted with planning mutations in TIR domains. The manuscript was written by B.H., S.B.C., V.H., and R.S. The study was supervised by V.H and R.S. All authors contributed to editing the manuscript and approved the final version.

## Supplementary figures

**Figure S1.**
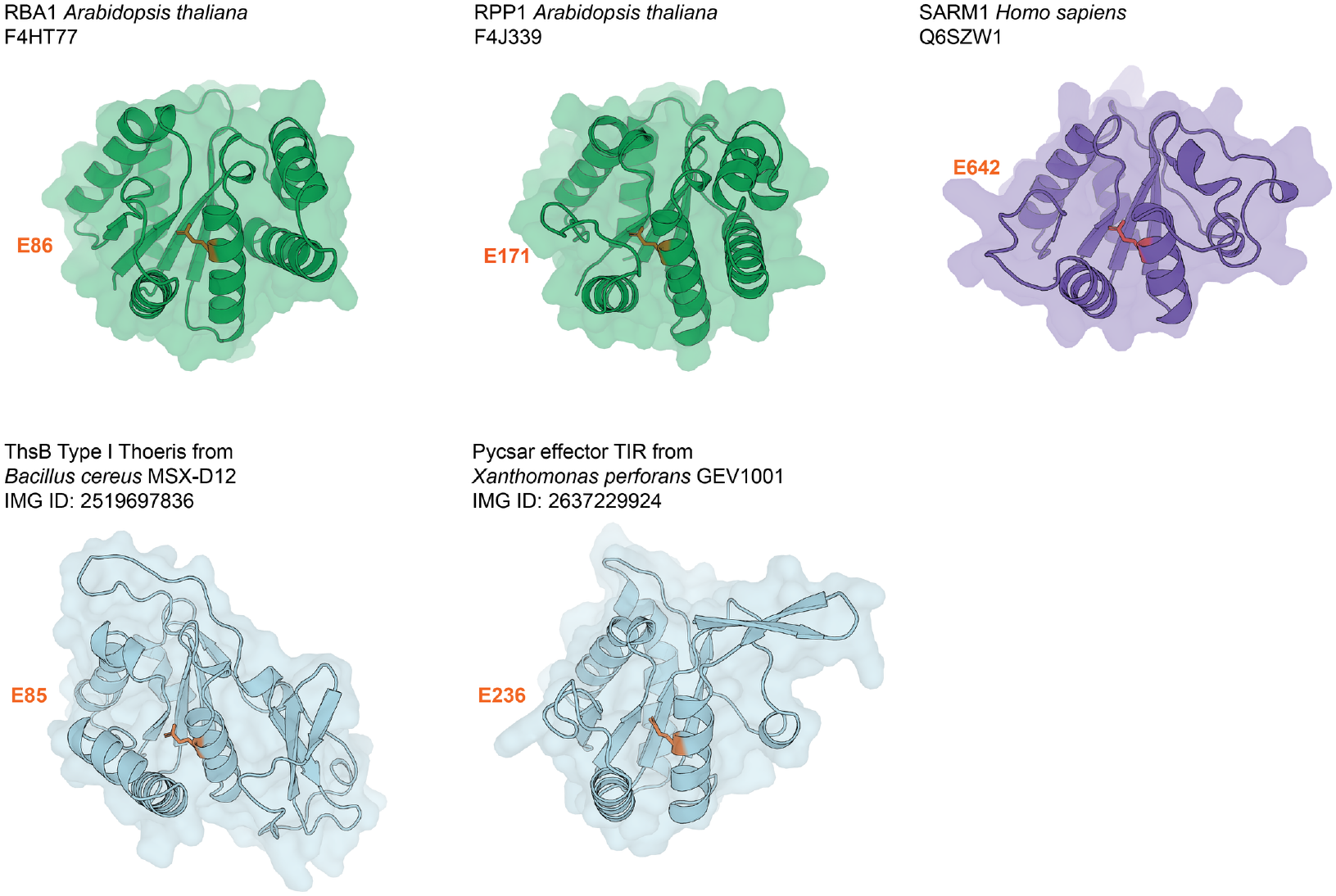
Predicted structures of catalytic TIR domains. AlphaFold3-predicted structures of known catalytic TIR domains from plants (green), human (violet), and bacteria (blue) with their catalytic glutamic acid residue highlighted in orange. UniProt/IMG ID and source species are indicated for each TIR domain. Related to Figure 1.

**Figure S2.**
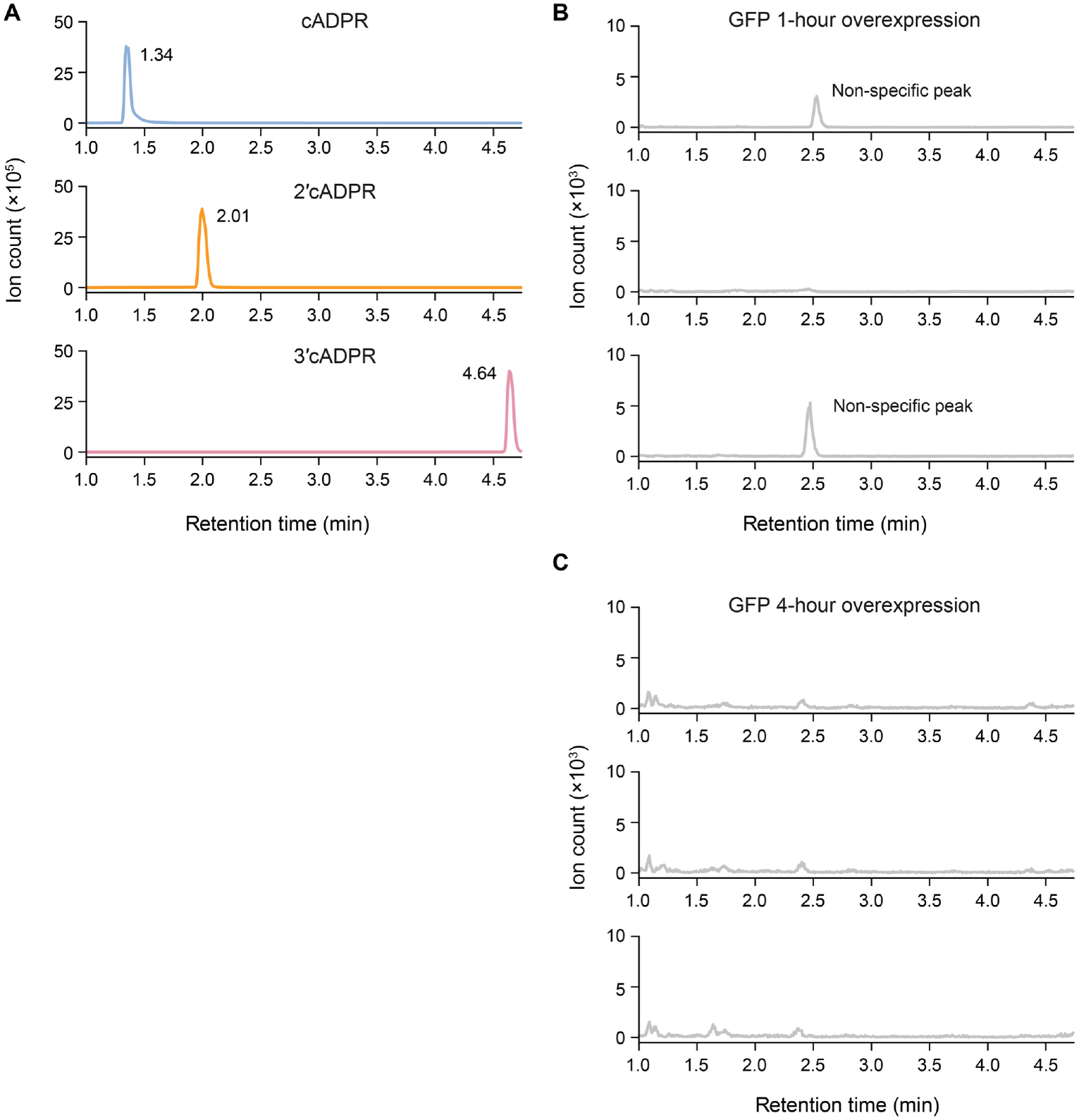
An LC-MS protocol for separating cADPR isomers. Extracted ion chromatograms (m/z = 540.053) of **(A)** cADPR, 2ʹcADPR, and 3ʹcADPR standards, showing distinct retention times, and of lysates of negative control cells expressing GFP following **(B)** 1 hour or **(C)** 4 hours of expression. Data in **(A)** are representative of three replicates; three independent replicates are shown in **(B)** and **(C)**. Related to Figures 1–3.

**Figure S3.**
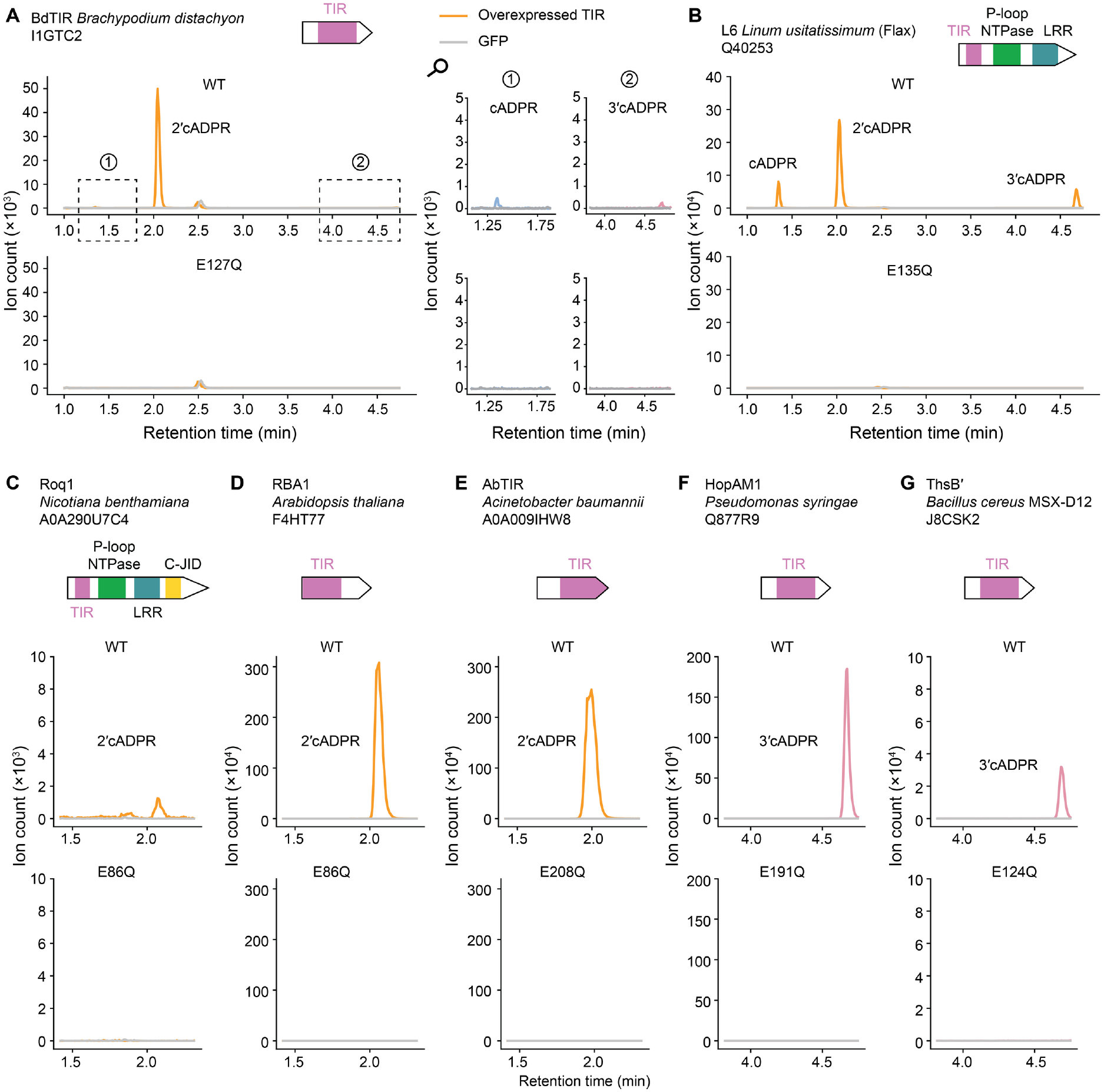
Known catalytic TIR domains produce cADPR isomers. Extracted ion chromatograms (m/z = 540.053) of *E. coli* cell lysates overexpressing TIR domains of previously reported proteins. Zoomed-in regions for BdTIR highlight minor peaks. UniProt IDs, source organisms, and protein domain architectures are indicated in each case. Data are representative of three replicates. The GFP control LC-MS data (1 hour induction) are also shown in Figures 2C, 2F, 2G, 3B, S4, S5, S11, S13, and S15. All three repeats of GFP control for both 1-hour and 4-hour induction are shown in Figure S2B and C. Related to Figure 1.

**Figure S4.**
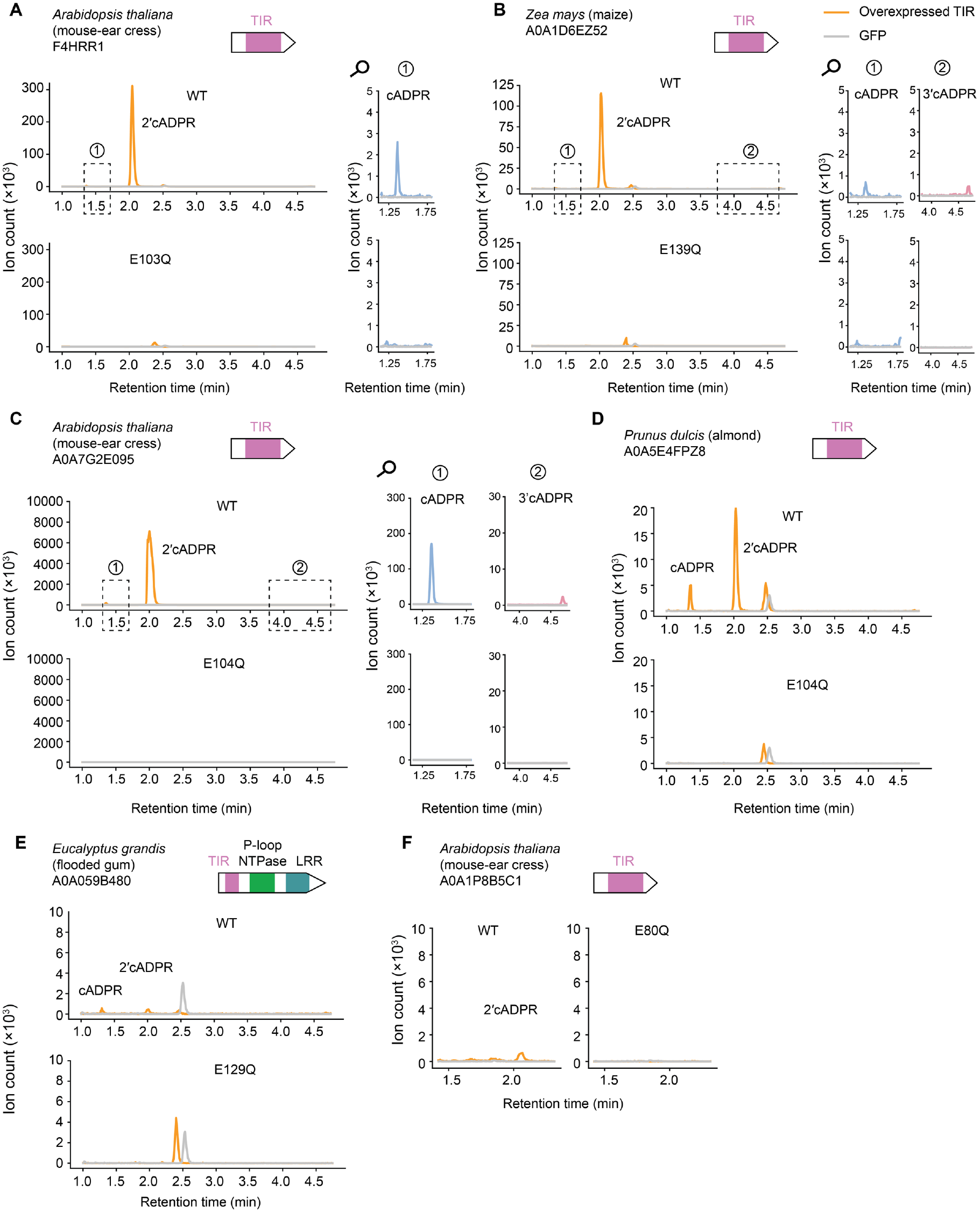
Plant TIR domains produce 2ʹcADPR as a main product. Extracted ion chromatograms (m/z = 540.053) of *E. coli* cell lysates expressing the TIR domains of plant proteins. UniProt IDs, source organisms, and protein domain architectures are indicated. Data are representatives of three replicates. The GFP control LC-MS data (1 hour induction) are also shown in Figures 2C, 2F, 2G, 3B, S3, S5, S11, S13, and S15. All three repeats of GFP control for both 1-hour and 4-hour induction are shown in Figure S2B and C. Related to Figure 1.

**Figure S5.**
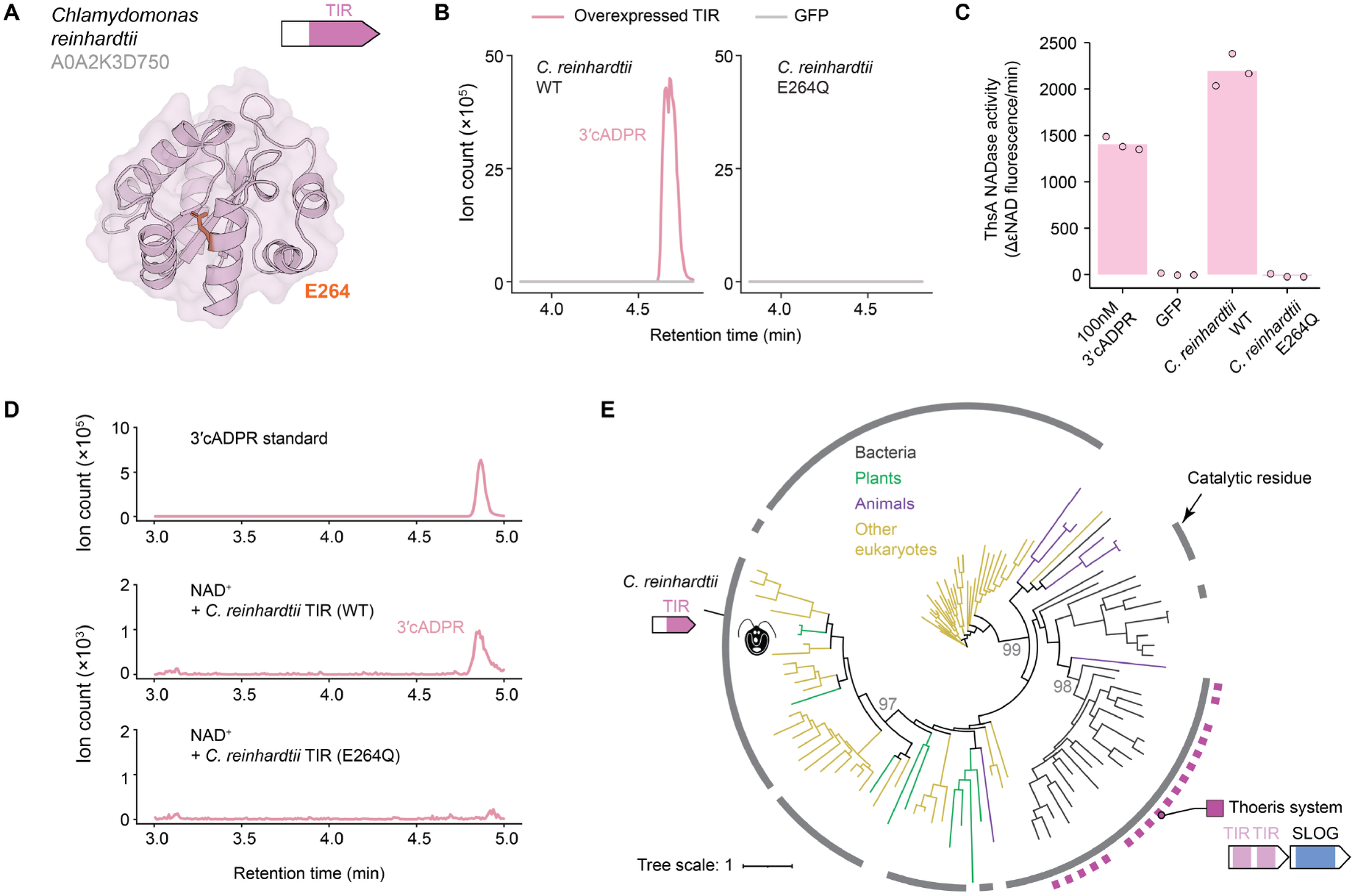
A green algal TIR domain synthesizes 3ʹcADPR. **(A)** AlphaFold3-predicted structure of the *Chlamydomonas reinhardtii* TIR domain, with the catalytic glutamic acid residue (E264) highlighted in orange. **(B)** Extracted ion chromatograms (m/z = 540.053) from lysates of *E. coli* cells expressing wild-type or catalytic mutant (E264Q) *C. reinhardtii* TIR domain for 1 hour. Data are representatives of three replicates. **(C)** NADase activity of purified *B. cereus* ThsA^5^ incubated with synthetic 3ʹcADPR (positive control) or with filtered lysates from *E. coli* cells expressing wild-type *C. reinhardtii* TIR or a catalytic mutant. Activity was measured using a nicotinamide 1,N6-ethenoadenine dinucleotide (εNAD) cleavage fluorescence assay. Bars represent the average of three experiments, with individual data points overlaid. **(D)** In vitro 3ʹcADPR production by purified *C. reinhardtii* TIR domain. Extracted ion chromatograms (m/z = 540.053) after incubation of NAD^+^ with or without the purified TIR domain. **(E)** Phylogenetic subtree of a TIR domain clade containing the *C. reinhardtii* TIR, extracted from a tree combining TIRs from eukaryotes and prokaryotes (Figure S7). Branch colors indicate the taxonomic group of the source organism. The outer ring denotes the presence (grey) of the predicted catalytic glutamic acid. Bacterial TIR domains encoded in operons with SLOG-domain genes are marked by violet rectangles. Bootstrap values are shown for major clades. The GFP control LC-MS data shown in panel B are also shown in Figures 2C, 2F, 2G, 3B, S3, S4, S11, S13, and S15. All three repeats of GFP control for both 1-hour and 4-hour induction are shown in Figure S2B and C. Related to Figure 1.

**Figure S6.**
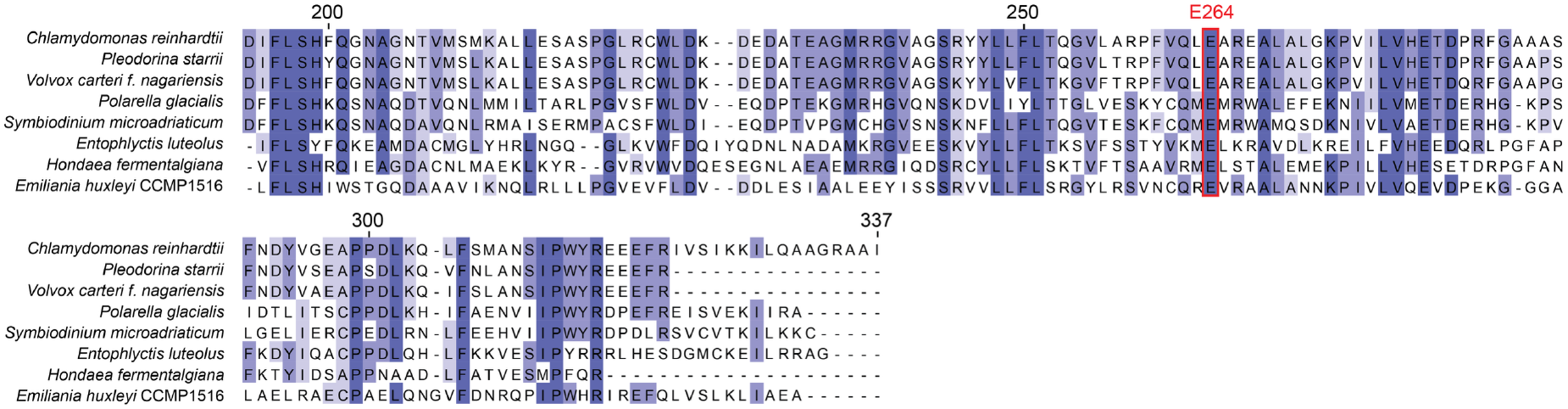
Diverse homologs of the *Chlamydomonas* TIR domain contain a conserved catalytic glutamic acid residue. Multiple sequence alignment of *C. reinhardtii* TIR domain homologs, with conserved residues highlighted in purple and catalytic residue shown in red. Source species are shown on the left. Related to Figure 1.

**Figure S7.**
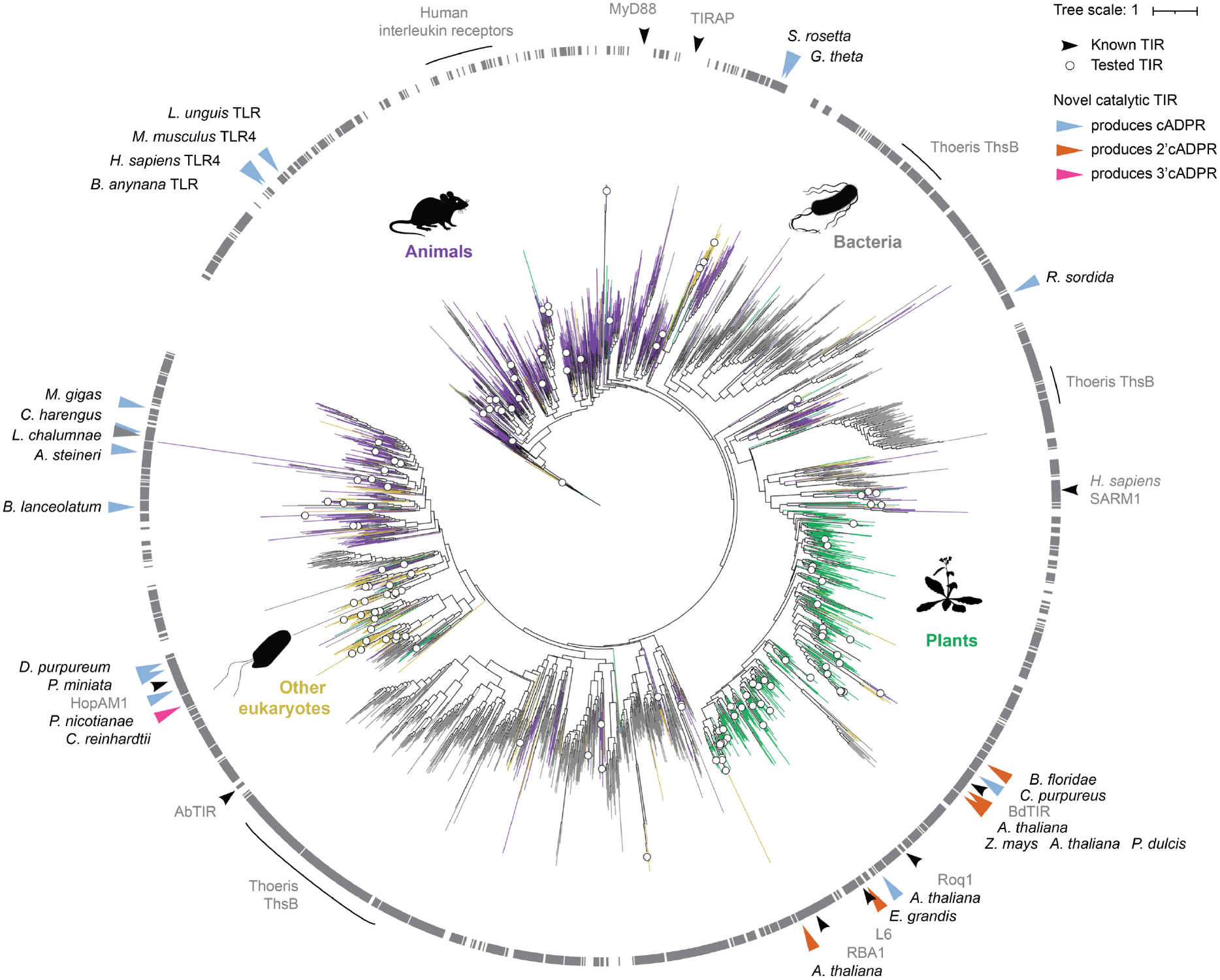
A phylogenetic tree of eukaryotic and prokaryotic TIR domains. A maximum-likelihood phylogenetic tree of ∼1,500 eukaryotic and 900 bacterial representative TIR domains. Branch colors indicate the taxonomic group of the source organism. The outer ring denotes the presence (grey) or absence of the conserved catalytic glutamic acid residue. White circles mark the 154 TIR domains experimentally tested in this study. TIR domains with detected catalytic activity are indicated on the tree periphery: black labels denote TIRs characterized in this study, colored by their major product (blue, cADPR; orange, 2ʹcADPR; pink, 3ʹcADPR). A TIR from *L. chalumnae* depleted NAD^+^ without generating a detectable cyclic ADPR product. Previously characterized TIRs are indicated in grey font. Related to Figure 1.

**Figure S8.**
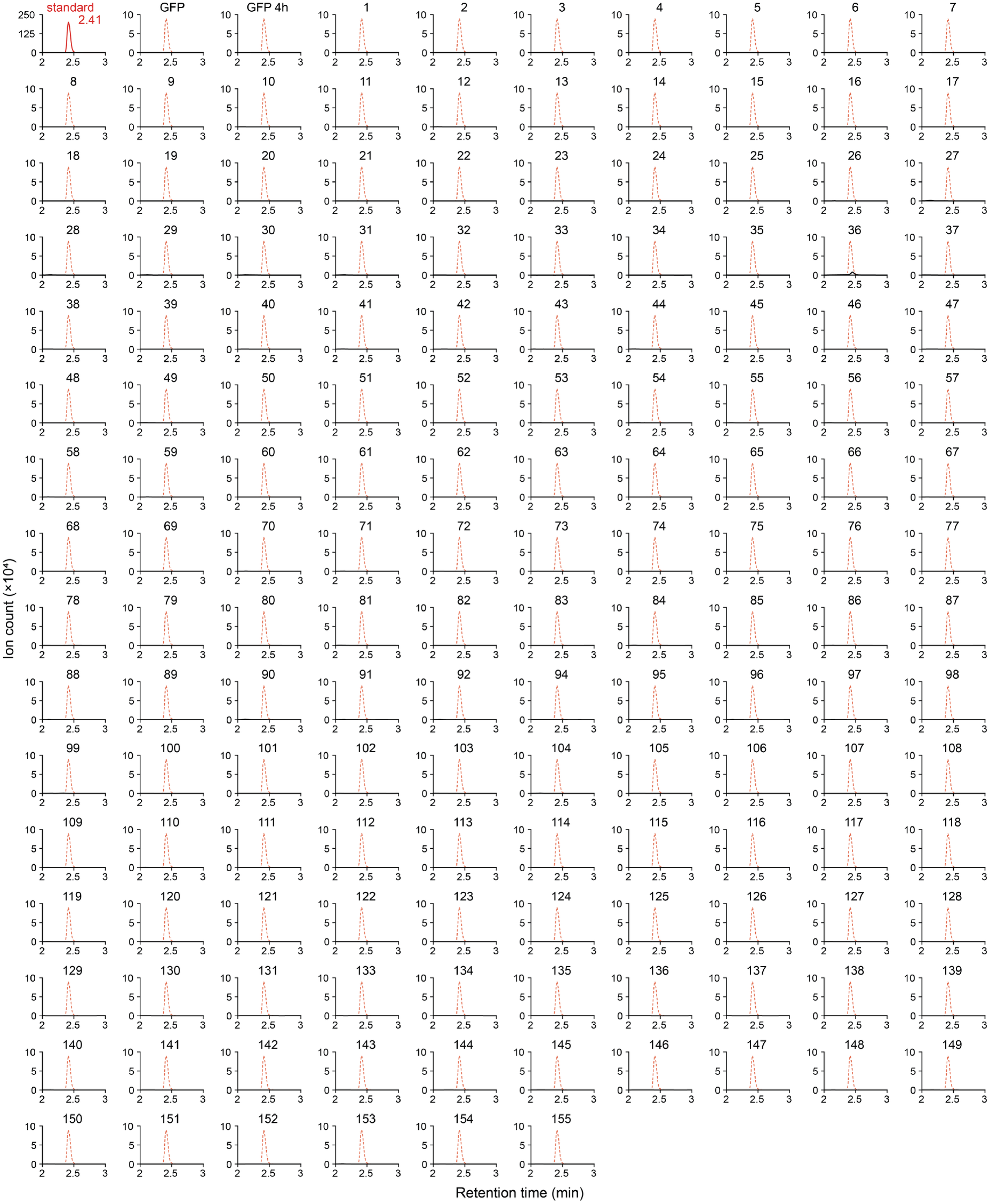
TIR domains screened here do not produce detectable His-ADPR. Extracted ion chromatograms (m/z = 695.126) of the His-ADPR standard (top left; retention time of the peak maximum indicated), of lysates of negative control cells expressing GFP following 1 hour (GFP) or 4 hours (GFP 4h) of expression induction, and of lysates of cells expressing each of the tested TIR-domain proteins, numbered as in Table S4. TIRs 19, 83, 122, 129, 139, 144, and 146 were sampled after 4 hours of expression. The dashed red line in each sample panel shows the His-ADPR standard chromatogram, scaled to the panel height, to mark its retention time. All samples except for the chemical standard are shown on the same scale (ion count x10^4^). One replicate is shown per sample. Related to Figure 1.

**Figure S9.**
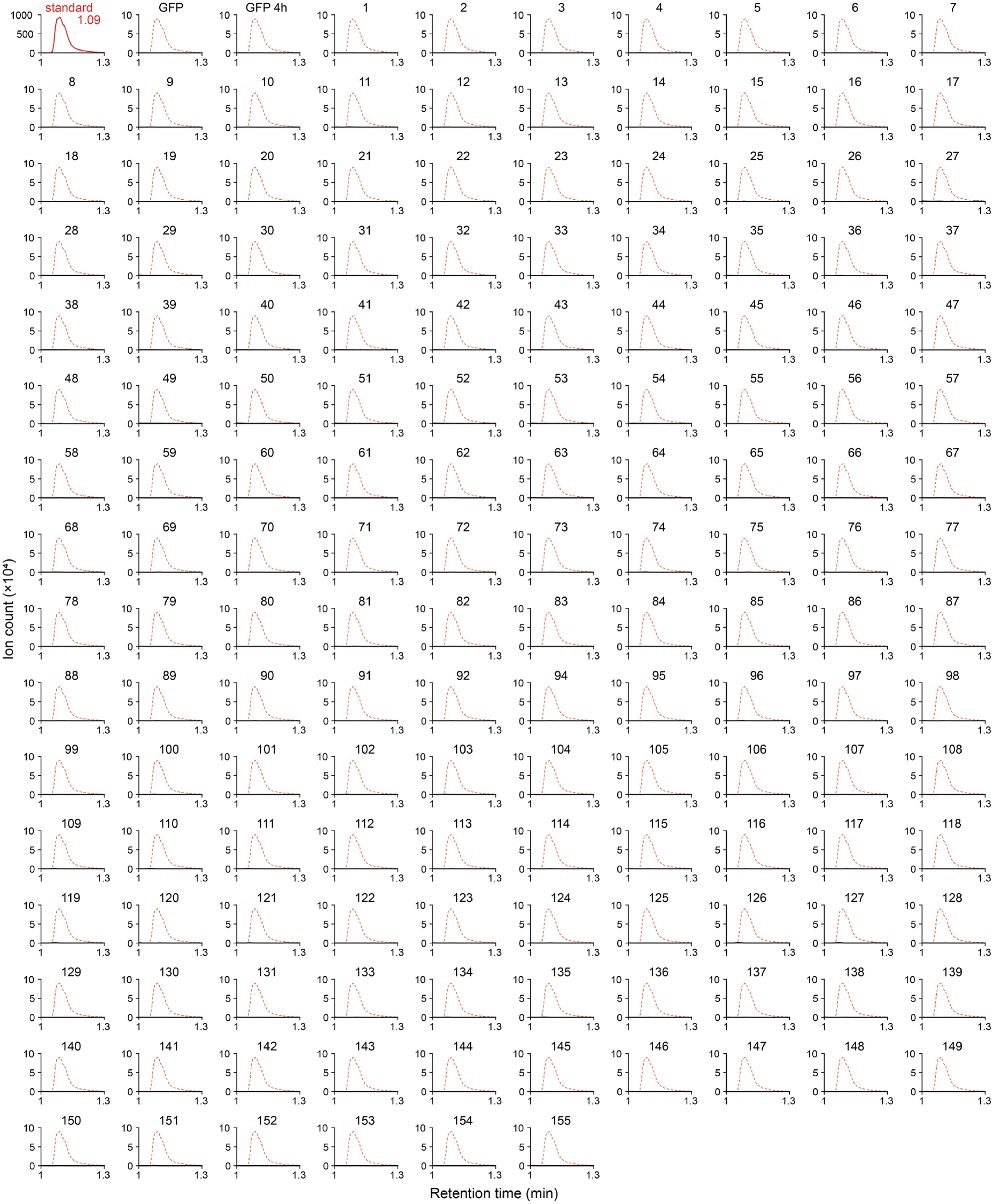
TIR domains screened here do not produce detectable pRib-AMP. Extracted ion chromatograms (m/z = 558.062) of the pRib-AMP standard (top left; retention time of the peak maximum indicated), of lysates of negative control cells expressing GFP following 1 hour (GFP) or 4 hours (GFP 4h) of expression induction, and of lysates of cells expressing each of the tested TIR-domain proteins, numbered as in Table S4. TIRs 19, 83, 122, 129, 139, 144, and 146 were sampled after 4 hours of expression. The dashed red line in each sample panel shows the pRib-AMP standard chromatogram, scaled to the panel height, to mark its retention time. All samples except for the chemical standard are shown on the same scale (ion count x10^4^). One replicate is shown per sample. Related to Figure 1.

**Figure S10.**
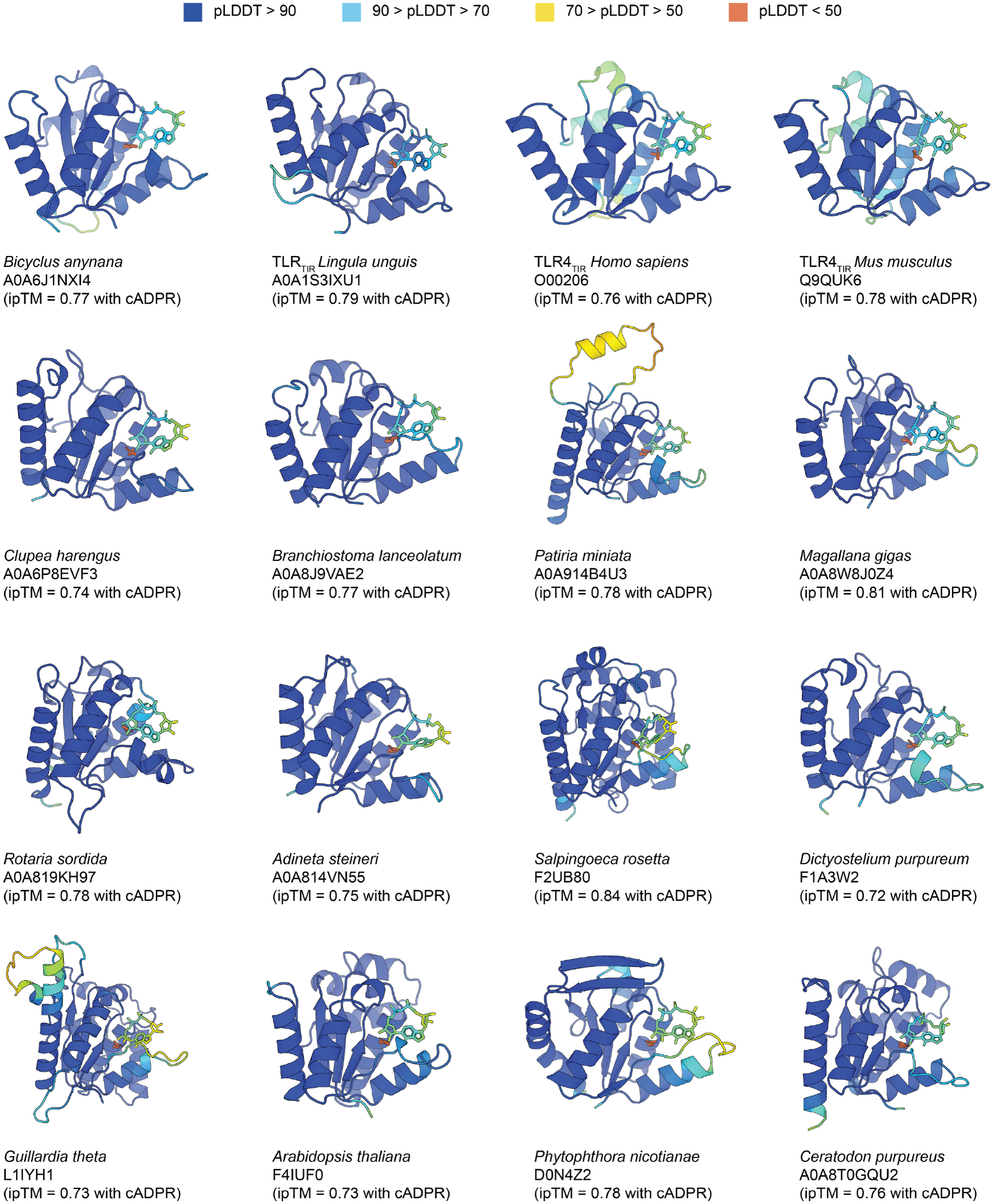
AlphaFold3 models of cADPR-producing TIR domains in complex with cADPR. Each panel shows a single TIR domain modeled together with cADPR. Proteins and cADPR are colored by their pLDDT score according to the legend. The conserved catalytic glutamate is shown in orange. Species names and UniProt ID are indicated below each panel, along with the ipTM value for the model. Related to Figures 2-3.

**Figure S11.**
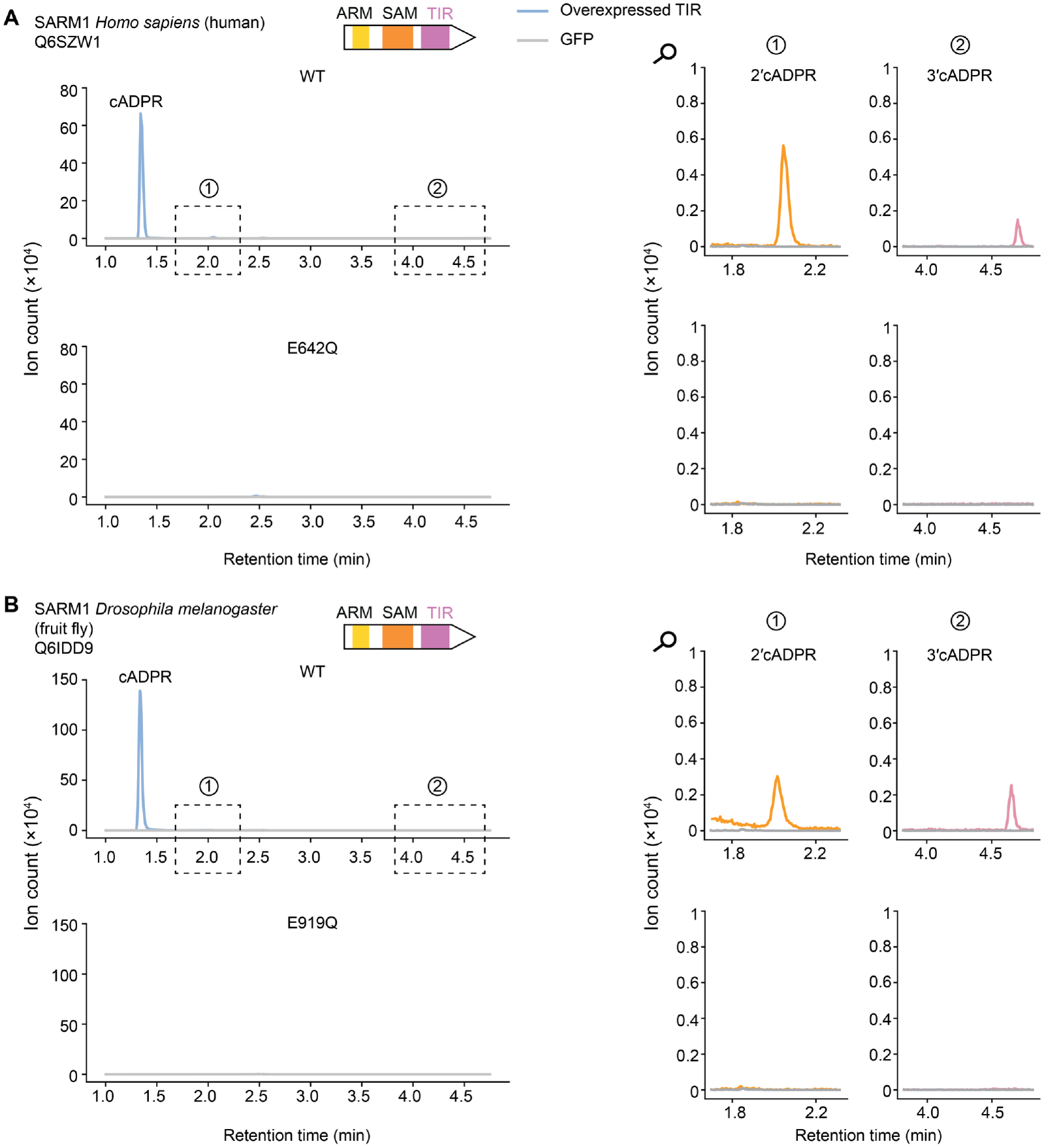
The SARM1 TIR domain produces cADPR as a major product. Extracted ion chromatograms (m/z = 540.053) of *E. coli* cell lysates overexpressing TIR domains of human and *Drosophila* SARM1. Zoomed-in regions highlight minor peaks. UniProt IDs and protein domain architectures are indicated. Data are representatives of three replicates. The GFP control LC-MS data (1 hour induction) are also shown in Figures 2C, 2F, 2G, 3B, S3, S4, S5, S13, and S15. All three repeats of GFP control for both 1-hour and 4-hour induction are shown in Figure S2B and C. Related to Figure 3.

**Figure S12.**
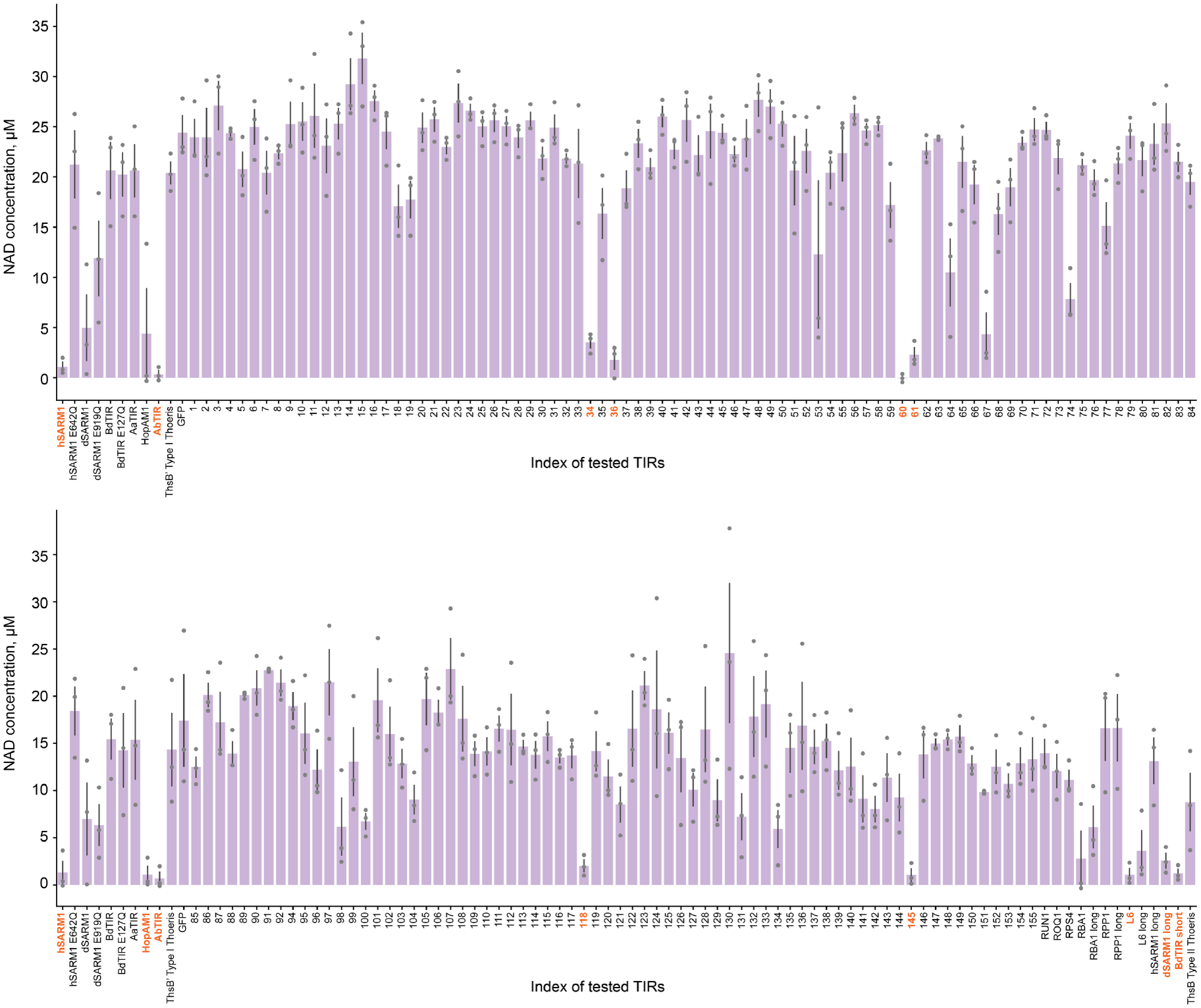
Concentration of NAD^+^ in bacterial cell lysates after overnight expression of TIR domains in this study. TIR domains were expressed overnight in bacteria, and the concentration of NAD^+^ was then measured in a 96-well plate format (Methods). Bars show the average of three repeats with individual data points overlayed. TIR domains that depleted most of the cellular NAD^+^ are highlighted in orange. The list of source organisms corresponding to TIR numbers is provided in Table S4. Related to Figure 3.

**Figure S13.**
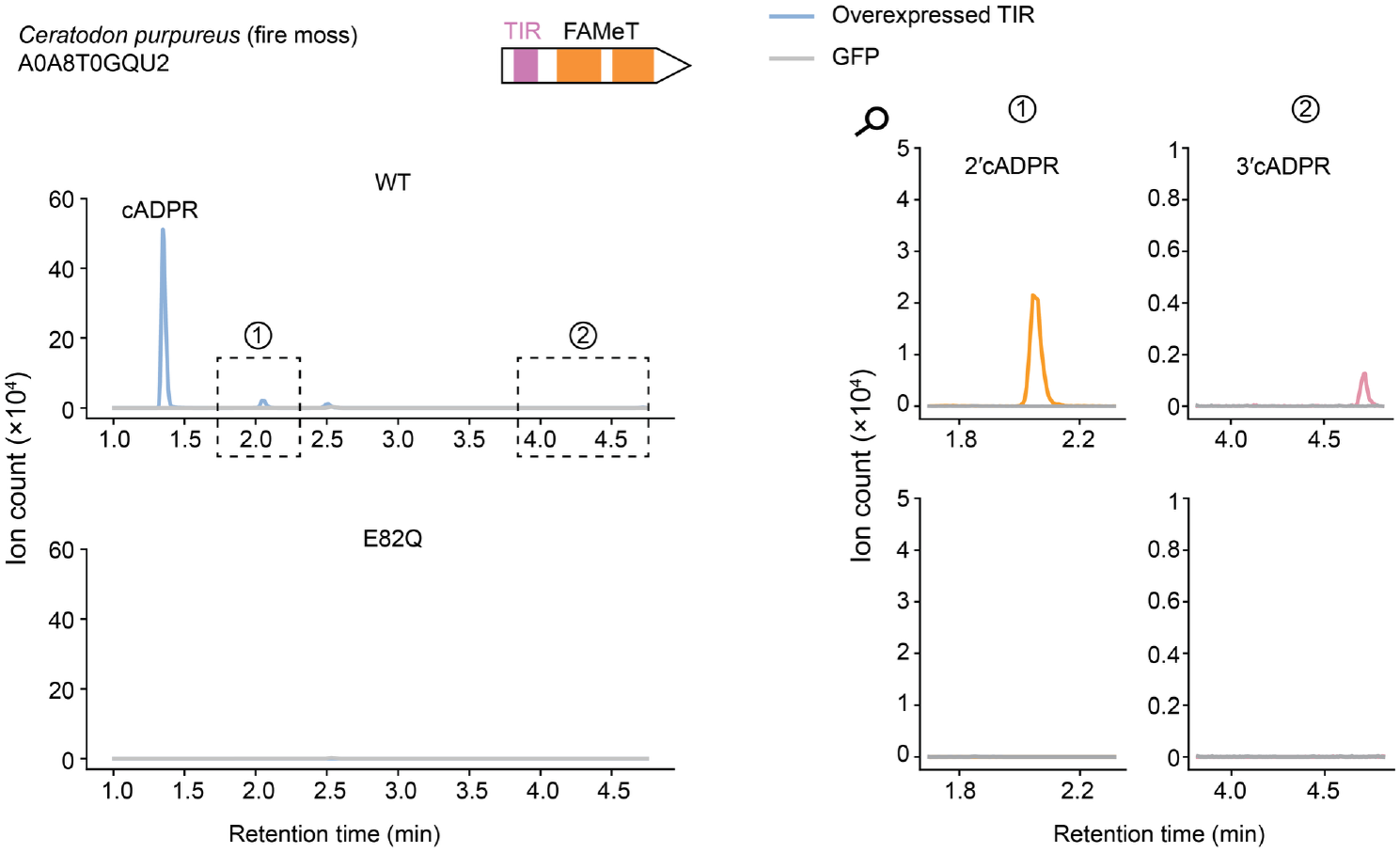
Activity of *Ceratodon* TIR domain. Extracted ion chromatograms (m/z = 540.053) of *E. coli* cell lysates overexpressing a TIR domain from *Ceratodon purpureus*. Zoomed-in regions highlight minor 2ʹcADPR and 3ʹcADPR peaks. UniProt ID and protein domain architectures are indicated. Data are representatives of three replicates. The GFP control LC-MS data (1 hour induction) are also shown in Figures 2C, 2F, 2G, 3B, S3, S4, S5, S11, and S15. All three repeats of GFP control for both 1-hour and 4-hour induction are shown in Figure S2B and C. Related to Figure 3.

**Fig. S14.**
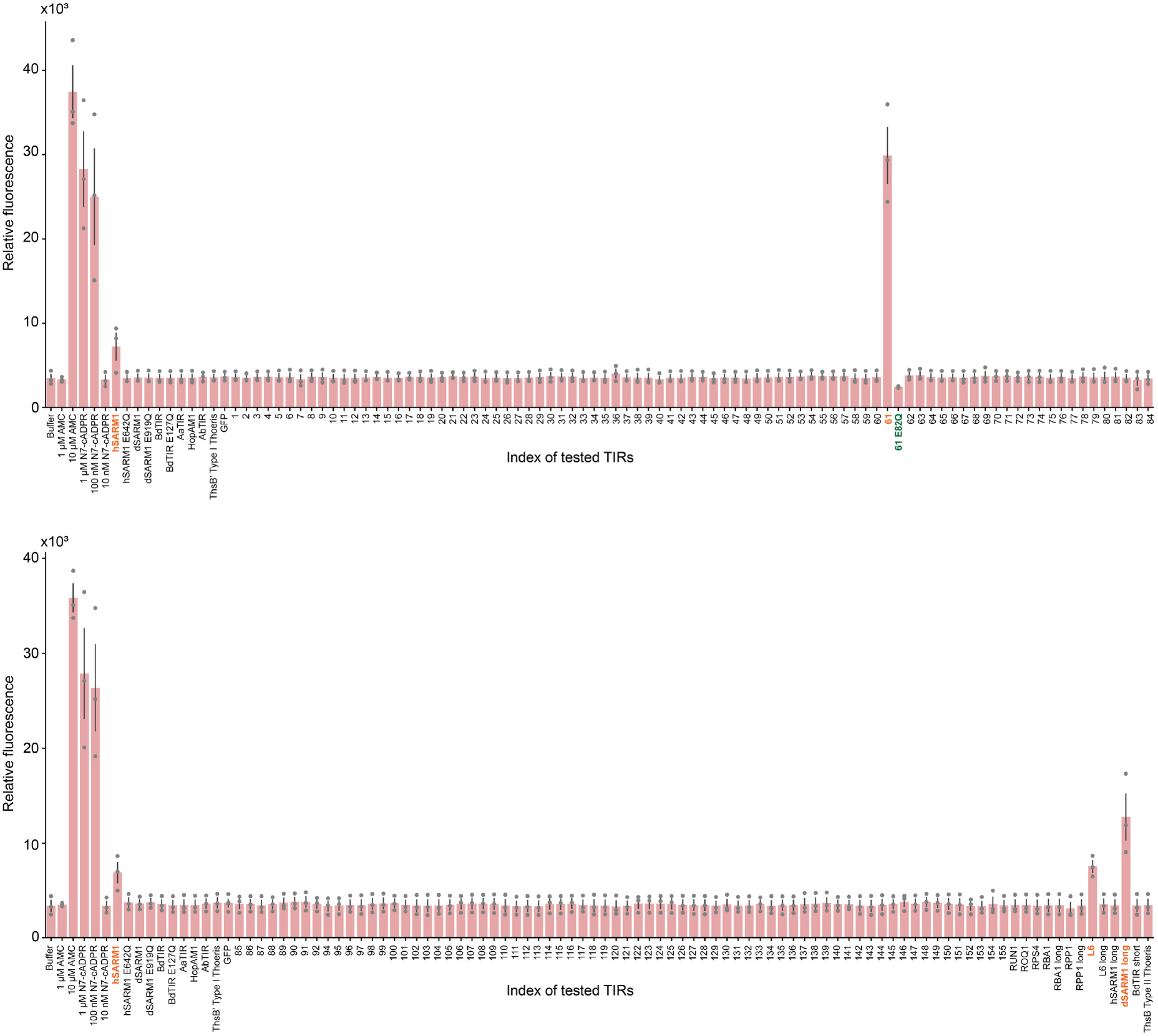
Measurement of N7-cADPR using a caspase activation assay. Activation of the caspase-like effector of the type IV Thoeris system, which is activated upon N7-cADPR binding^9^, was measured for lysates of cells expressing each TIR domain. TIR domains producing N7-cADPR in amounts sufficient to activate the effector are marked in orange, and their corresponding catalytic mutants, where included, in green. Bars show the average of three repeats with individual data points overlayed. The full list of source organisms corresponding to TIR numbers is provided in Table S4. Related to Figure 3.

**Figure S15.**
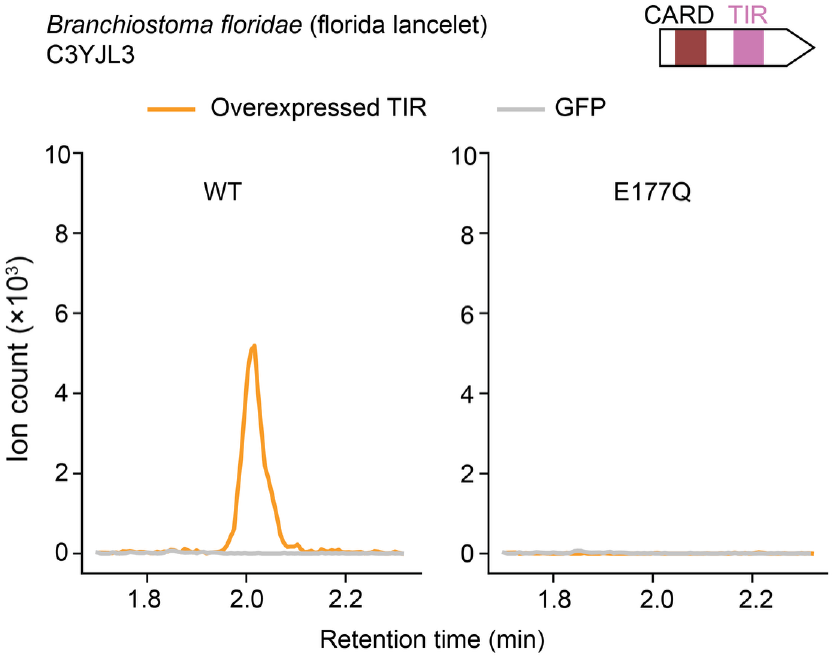
Lancelet TIR domain produces 2ʹcADPR. Extracted ion chromatograms (m/z = 540.053) of *E. coli* cell lysates overexpressing a TIR domain from a *Branchiostoma floridae* protein, with its UniProt ID and protein domain architecture indicated. Data are representatives of three replicates. The GFP control LC-MS data (1 hour induction) are also shown in Figures 2C, 2F, 2G, 3B, S3, S4, S5, S11, and S13. All three repeats of GFP control for both 1-hour and 4-hour induction are shown in Figure S2B and C. Related to Figure 3.

**Figure S16.**
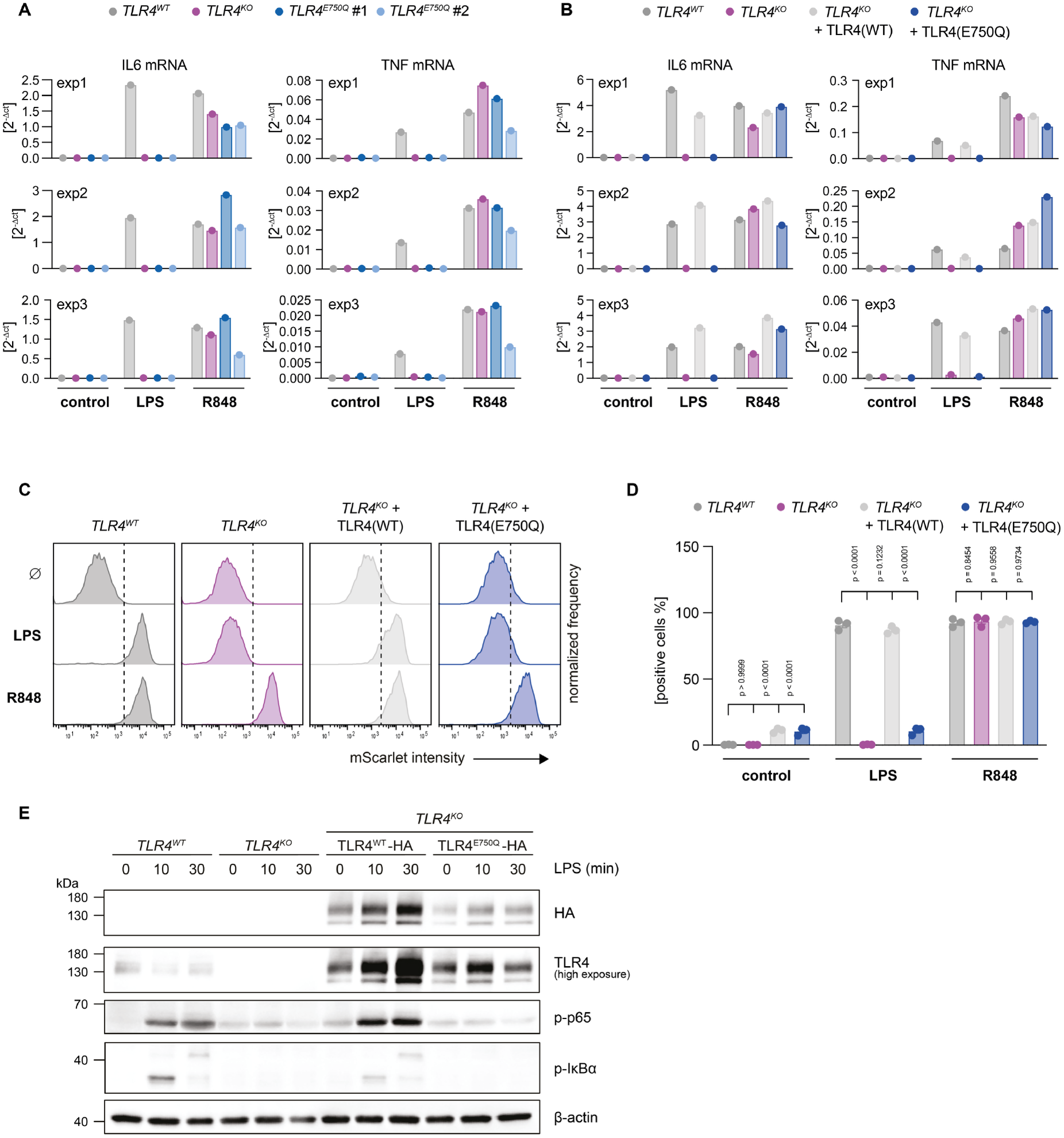
The TLR4 E750Q substitution impairs LPS-induced inflammatory signaling in human macrophages. **(A)** WT cells, TLR4 KO cells, and two independent TLR4 E750Q knock-in clones were left untreated or stimulated with LPS or R848 for 16 hours. Expression of *IL6* and *TNF* was analyzed by qPCR and is shown as 2^−ΔCt^ relative to *GAPDH*. Results from three independent experiments are shown separately. **(B)** WT cells, TLR4 KO cells, and TLR4 KO cells reconstituted with doxycycline-inducible WT or E750Q TLR4 were left untreated or stimulated with LPS or R848 for 16 hours. Expression of *IL6* and *TNF* was analyzed by qPCR and is shown as 2^−ΔCt^ relative to *GAPDH*. Results from three independent experiments are shown separately. **(C)** CEBRA cells of the indicated genotypes were left untreated or stimulated with LPS or R848, and mScarlet reporter expression was measured by flow cytometry 16 hours after stimulation. Histograms from one representative experiment are shown. Dashed vertical lines indicate the threshold used to identify mScarlet-positive cells. **(D)** Quantification of the experiment described in (C). Data are presented as mean ± SEM of three independent experiments. Statistical significance was determined by two-way ANOVA followed by Dunnett’s multiple-comparisons test. **(E)** *TLR4* WT or *TLR4* knockout CEBRA cells reconstituted with doxycycline-inducible HA-tagged WT or E750Q TLR4 were stimulated with LPS for the indicated times. NF-κB pathway activation was assessed by Western blot analysis of phosphorylated p65 and IκBα. TLR4 expression was assessed by blotting for TLR4 or the HA tag, as indicated, and β-actin served as a loading control. One representative experiment of three independent experiments is shown. Related to Figure 4.

**Figure S17.**
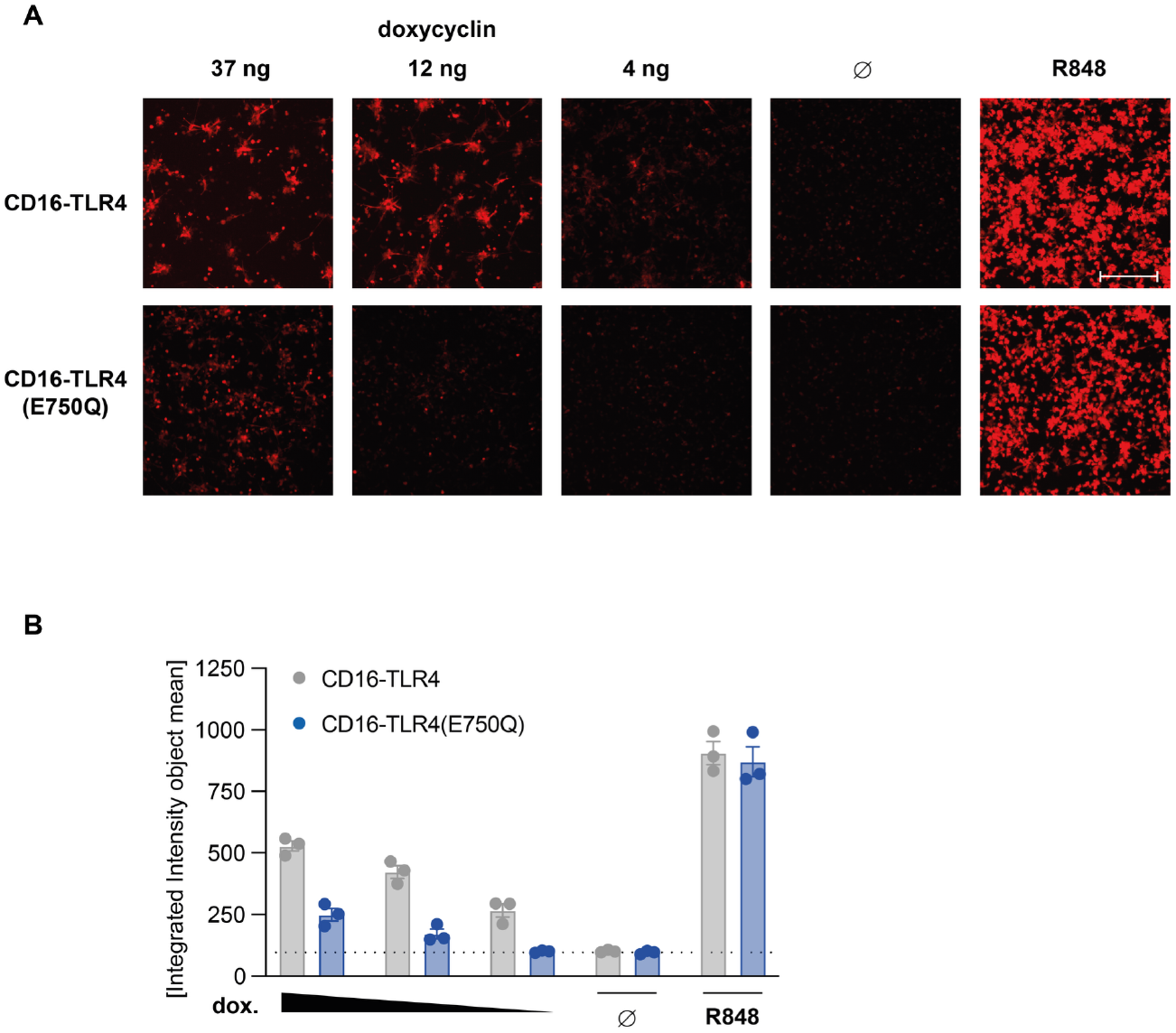
The TLR4 catalytic residue is required for signaling by the CD16–TLR4-TIR chimeric receptor. **(A)** CEBRA cells expressing doxycycline-inducible WT or E750Q CD16–TLR4-TIR constructs were treated with 37, 12, or 4 ng/ml doxycycline, left untreated, or stimulated with R848. At 24 hours after induction, *IL1B*-mScarlet reporter expression was analyzed by Incucyte imaging. Representative fields from one of three independent experiments are shown. Scale bar, 200 µm. **(B)** Quantification of *IL1B*-mScarlet reporter expression from three independent experiments performed as in (A). Reporter activity was quantified as mean integrated intensity per object. Data are presented as mean ± SEM. The dotted horizontal line indicates the background fluorescence. Related to Figure 5.

**Figure S18.**
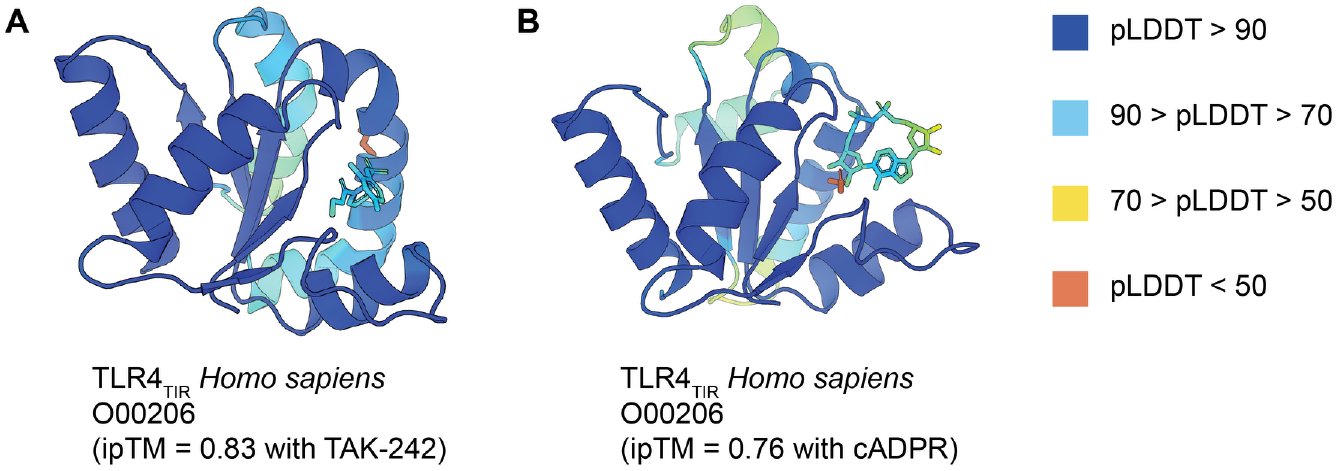
AlphaFold3 models of TLR4 TIR domain in complex with diverse ligands. Each panel shows a single TIR domain modeled together with **(A)** TAK-242 (resatorvid), or **(B)** cADPR. Proteins and ligands are colored by their pLDDT score according to the legend. The conserved catalytic glutamate is shown in orange. UniProt ID and the ipTM value for the model are indicated below each panel. Related to Figure 2.

## Star Methods

### Key resources table

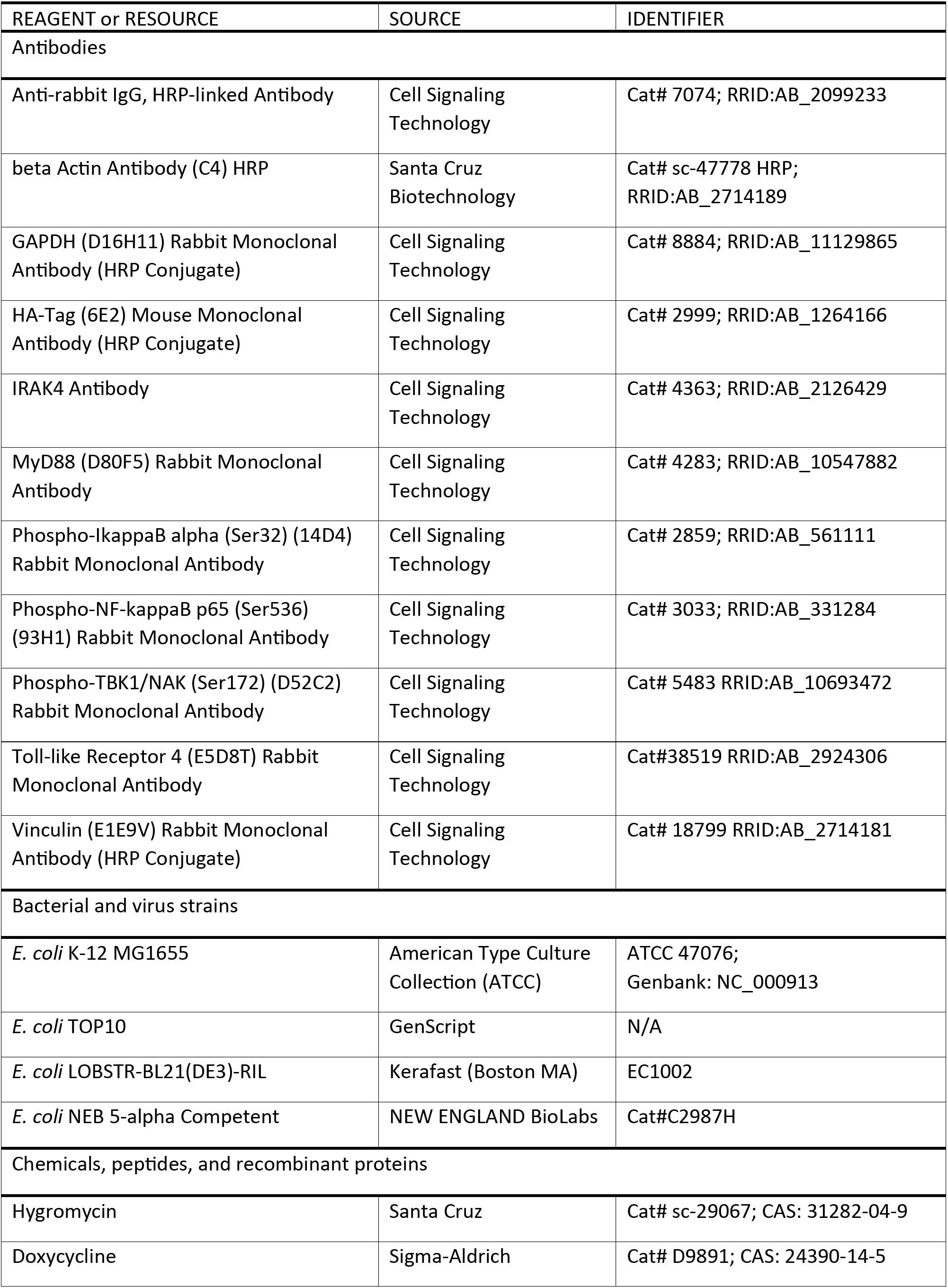
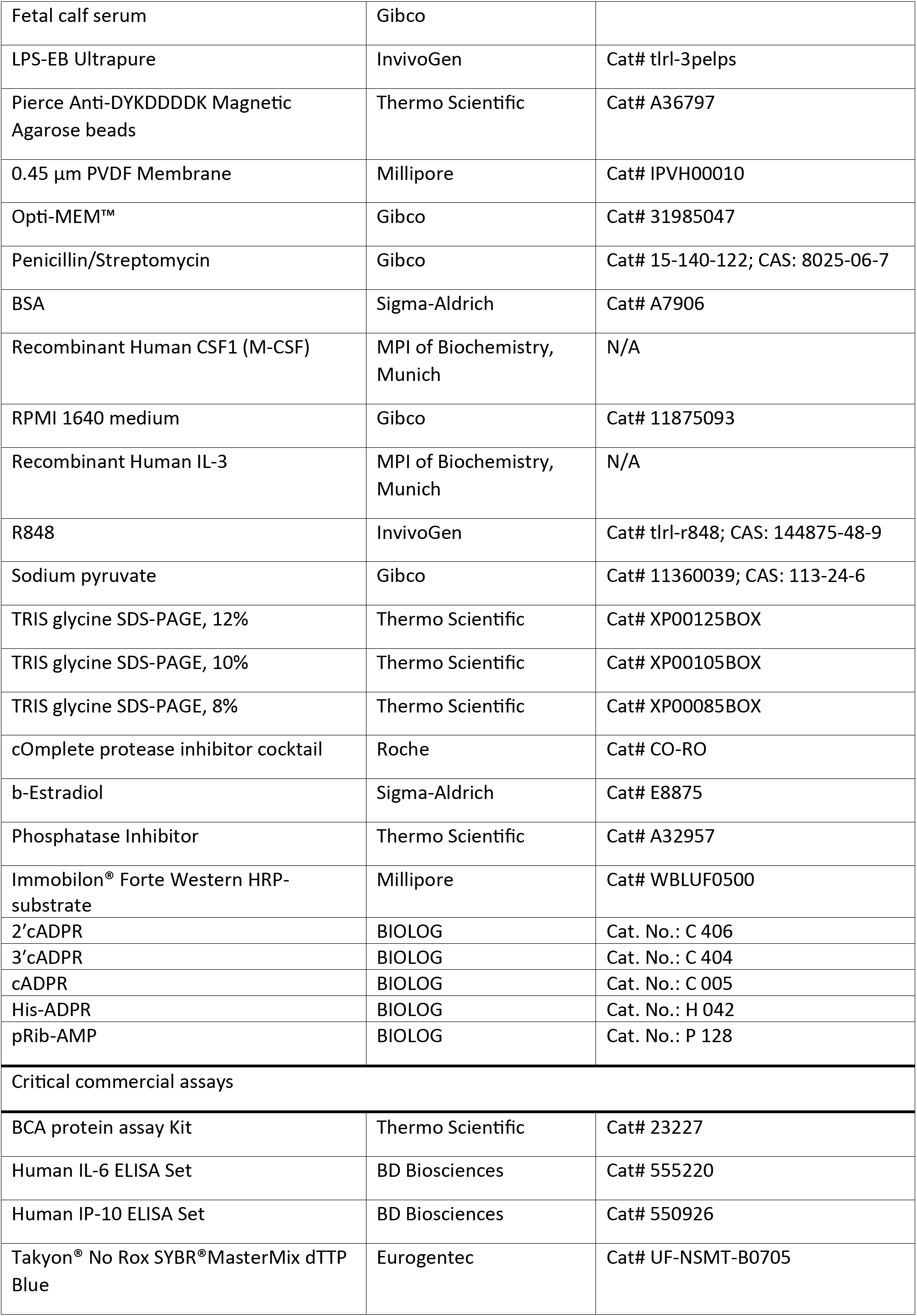
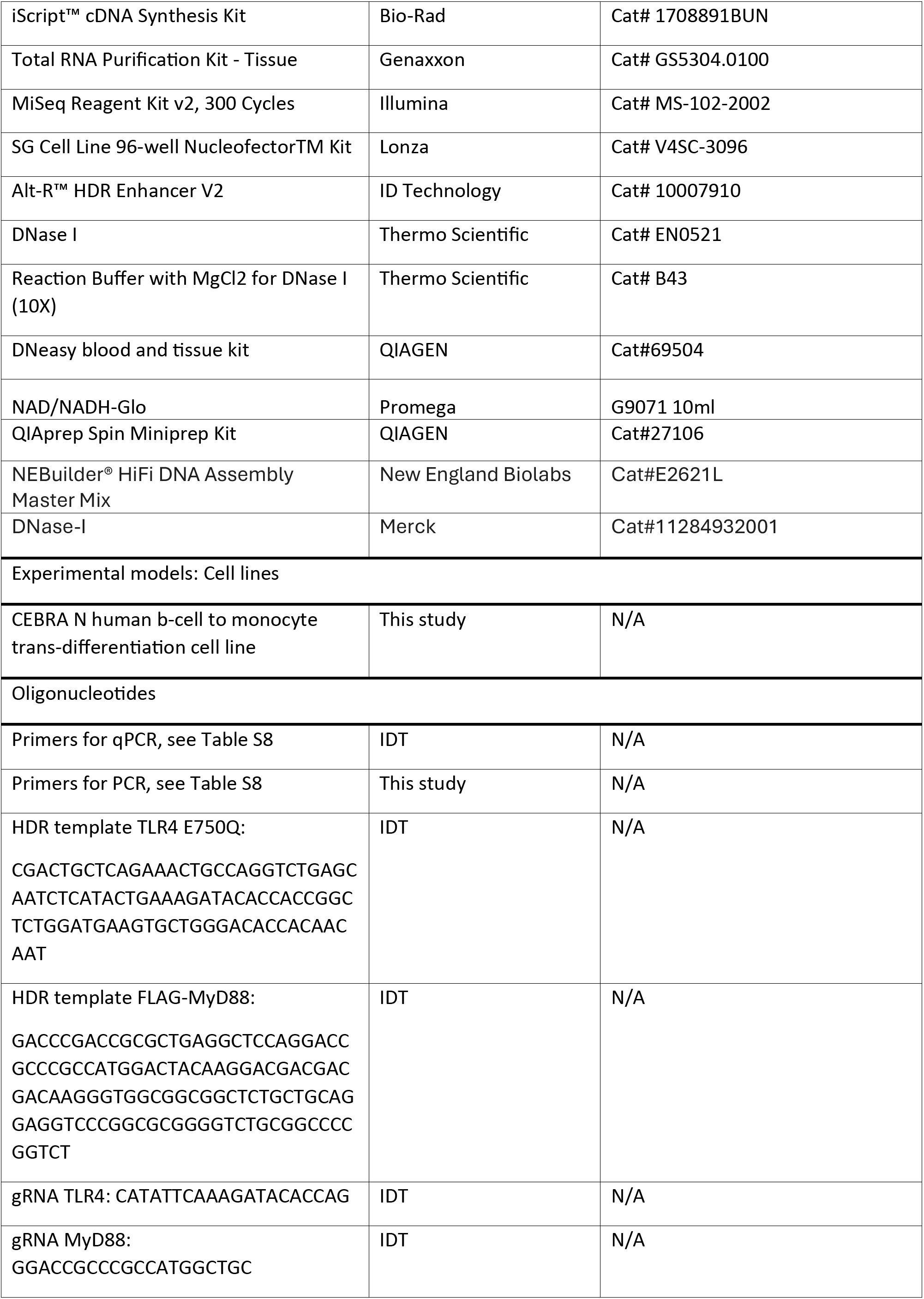
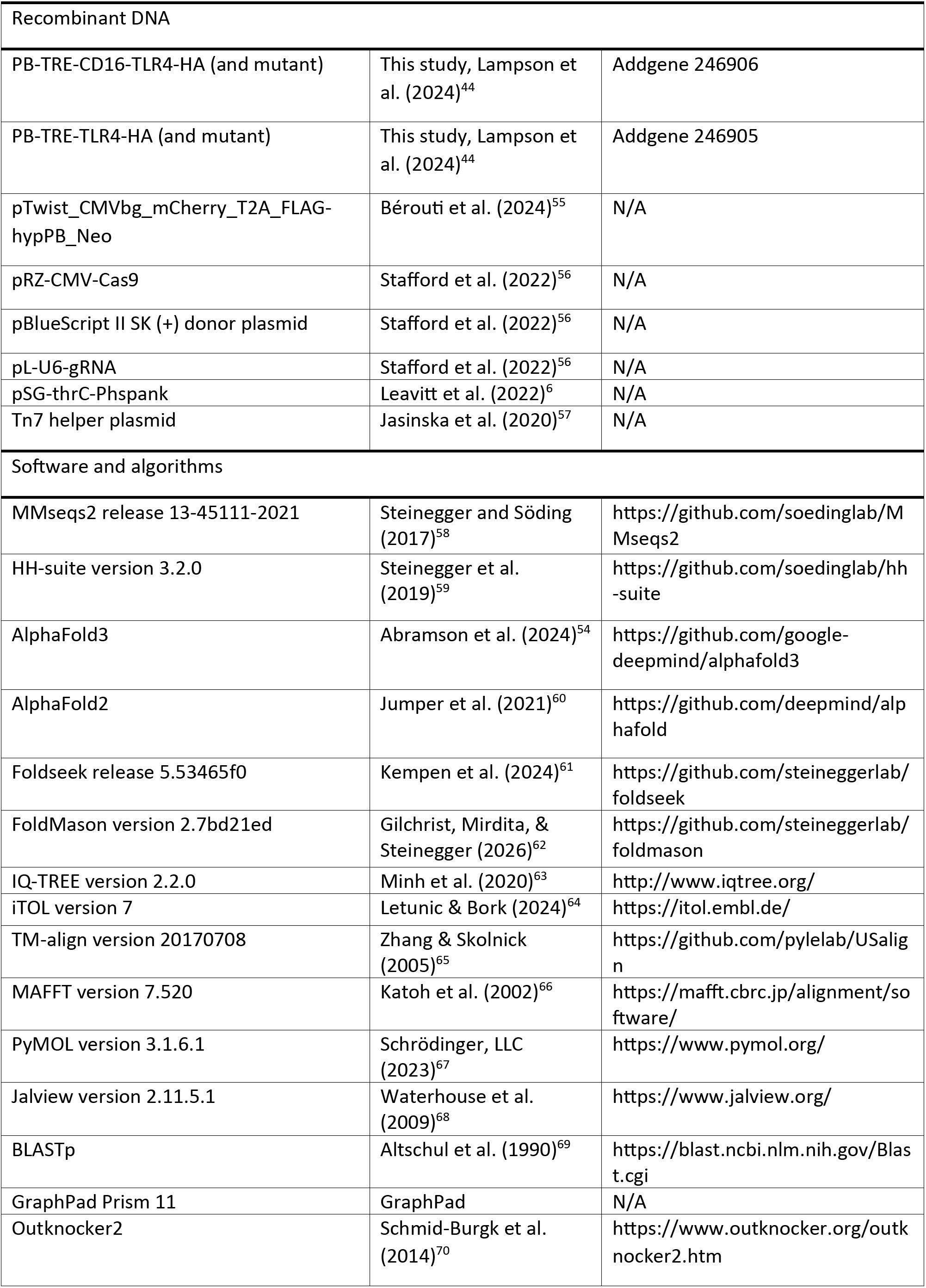
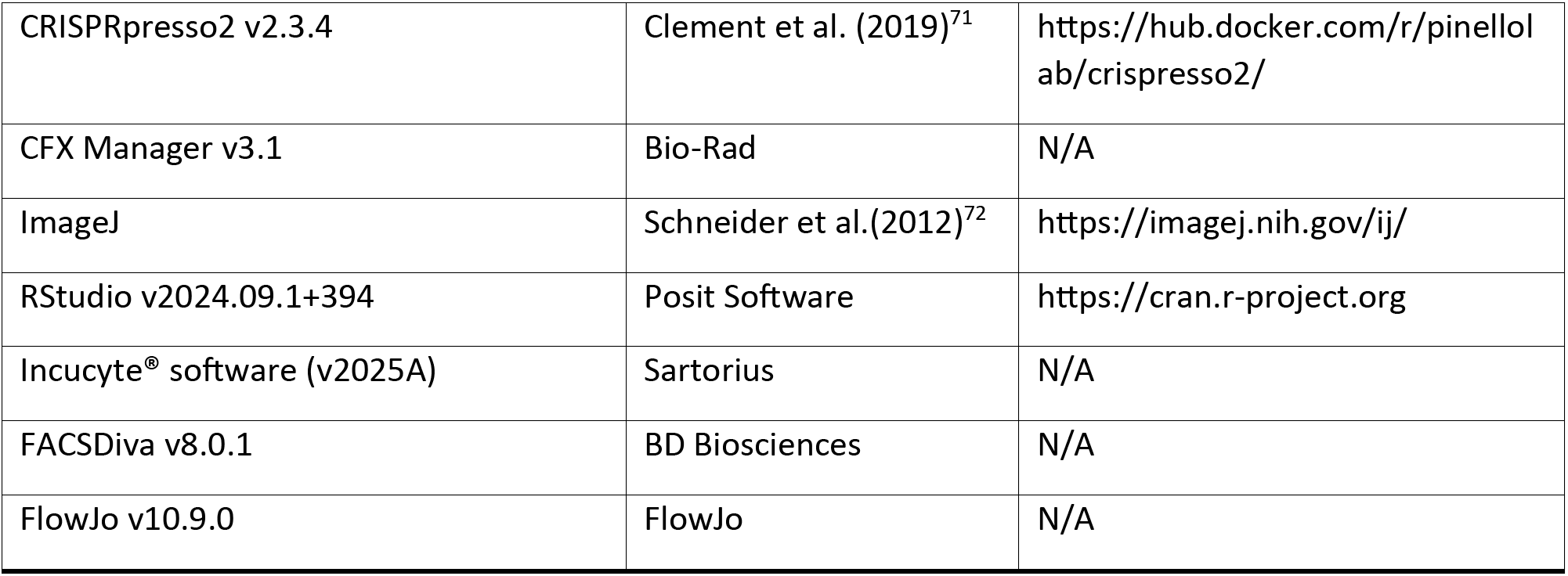

## Experimental Model and study participant details

### Bacterial strains

All *E. coli* strains were grown in MMB media (lysogeny broth (LB) supplemented with 0.1 mM MnCl2 and 5 mM MgCl2) at 37°C with 200 rpm shaking. Ampicillin 100 μg/mL or kanamycin 50 μg/mL were added when necessary for plasmid maintenance. The following strains of *E. coli*: TOP10 (GenScript) was used for TIR overexpression, DH5a (NEB) was used for cloning, LOBSTR-BL21(DE3)-RIL for protein purification, and K-12 MG1655 for toxicity experiments with bacterial Thoeris systems. All chemicals were obtained from Sigma-Aldrich and standard molecules from BIOLOG unless stated otherwise.

Thoeris type VII from the *Alteromonadales* metagenomic scaffold^8^ was integrated into the *E. coli* MG1655 genome using the Tn7 integration plasmid as previously described^73^ (spectinomycin resistance, induced by anhydrotetracycline, Sigma cat.37919)^57^. A plasmid expressing RFP was integrated as a negative control.

All plasmids were synthesized by GenScript Corporation or Twist Bioscience. TIR domain genes and GFP as a control were codon optimized for expression in *E. coli* and synthesized by GenScript. These constructs were subsequently cloned into the pSG-thrC-Phspank^6^ plasmid under the IPTG-inducible promoter. Transformations into *E. coli* were performed using standard electroporation or TSS^74^. All transformed strains were validated by a plasmid sequencing service (Plasmidsaurus).

### Cell model system

The CEBRA (**C/EB**Pα-E**R a**ctivated transdifferentiation system) cell line (in the following referred to as CEBRA) was generated by the stable integration of a fusion construct of *C/EBPα* and the ligand-binding domain of the estrogen receptor (*C/EBPα-ESR1LBD*) into RCH-ACV cells using the PiggyBac transposon system^75^. RCH-ACV is a lymphoblastic leukemia cell line with chromosome translocation 1; 19 and trisomy 8. Controlled nuclear translocation of the C/EBPα-ESR1LBD fusion protein, induced by β-estradiol in the presence of interleukin-3 and macrophage colony-stimulating factor, induces the conversion of B-lineage cells to monocytes/macrophages, while ensuring cell viability. The cell line carries a nourseothricin-resistance cassette.

### Cell culture

CEBRA cells were cultured in RPMI1640 medium, supplemented with 10% heat-inactivated fetal calf serum (FCS), 100 U/mL penicillin, 100 µg/mL streptomycin, and 1 mM sodium pyruvate. Cells were cultured in a humidified incubator at 37 °C with 5% CO_2_.

For transdifferentiation of CEBRA cells into macrophages, cells were seeded in culture medium supplemented with the cytokines M-CSF (10 ng/mL), IL-3 (10 ng/mL), and b-estradiol (100 nM) for 5 days. Cells were seeded in 96-well plates (80,000 cells/well) or in non-tissue culture-treated 6-well plates or 10-cm dishes (600,000 cells/mL). Cells differentiated in 6-well plates and 10-cm dishes were re-seeded in 96-well plates (100,000 cells/well) on day 5 for downstream experiments, unless otherwise indicated.

### Cell stimulation

Differentiated CEBRA cells were treated with LPS, R848, and/or doxycycline at the concentrations and for the durations indicated in the respective figure legends. Cells and culture supernatants were collected at the indicated experimental endpoints for downstream analysis.

### Plasmid electroporation

CEBRA cells were seeded at a density of 200,000 cells/mL and cultured overnight. The following day, 2.5×10^6^ cells were resuspended in 250 µL Opti-MEM, mixed with plasmid DNA, and incubated for 20 min. Next, the cell suspension was transferred to a 4-mm electroporation cuvette, electroporated (265 V, 975 µF, 720 W) using a Gene Pulser (Bio-Rad), and immediately transferred to 6-well plates with pre-warmed medium.

### CRISPR/Cas9-mediated knockout generation

Knockouts in CEBRA cells were generated using ribonucleoprotein (RNP) delivery. RNPs were assembled by annealing 1:1 molar ratio of tracrRNA and crRNA at 95 °C for 5 min, followed by stepwise (–1 °C/35 sec) decrease to 22 °C. NLS-Cas9 (40 pmol) was added to each gRNA complex (100 pmol), incubated at room temperature for 20 min, and frozen at -80 °C. The cells were nucleofected using the SG Cell Line 96-well Nucleofector kit (Lonza). 500,000 cells were resuspended in 20 µL of SG Cell Line Nucleofector Solution, and the Cas9/RNP assembly was added. The cells were nucleofected on a 4D-Nucleofector (Lonza) using the program CM-130. Subsequently, the cells were transferred to pre-warmed culture medium. Cells were rested for 48 h, then seeded at limiting dilutions to obtain monoclonal populations. After three weeks, monoclones of CEBRA cells were picked and analyzed by MiSeq as described before^70^. Homozygous clones were validated using Outknocker2^70^ or CRISPRpresso2^71^. Functional validation of selected clones was conducted by Western Blotting and stimulation experiments.

### CRISPR/Cas9-mediated knock-in generation

Cas9 (43.4 pmol) and sgRNA (100 pmol) were complexed for 20 min at room temperature. Next, the single-stranded DNA donor (150 pmol) was added. Cells were nucleofected as described above and re-seeded in pre-warmed culture medium containing 1 µM HDR enhancer (IDT). After 24 h, the medium was exchanged to remove the HDR enhancer, and 2 days later, cells were seeded at limiting dilutions to obtain monoclonal populations. For point mutations, monoclones were analyzed by MiSeq as described above. For the endogenous tagging of *MyD88* with a FLAG-tag, monoclonal clones were analyzed by PCR and selected clones further analyzed by Sanger sequencing.

### Generation of the endogenous *IL1B*-T2A-mScarlet reporter cell line

The CEBRA IL1B^mScarlet^ reporter cell line was generated as previously described^56^. Briefly, cells were electroporated as described above using a pRZ-CMV-Cas9 plasmid encoding Cas9, a pBlueScript II SK (+) donor plasmid containing T2A-threonine-glycine-mScarlet cassette flanked by homology arms, and two pL-U6-gRNA plasmids encoding sgRNAs targeting sequences upstream and downstream of the *IL1B* stop codon (Cas9:gRNA:template equal to 4:4:1, 8 µg total DNA). The sgRNA recognition sites flanking the donor cassette enable Cas9-mediated release of the knock-in cassette, while the corresponding recognition sequences within the donor DNA were disrupted by synonymous mutations to prevent re-cleavage following integration. After electroporation, cells were rested for 48 h, then seeded at limiting dilutions to obtain monoclonal populations. After three weeks, monoclones of CEBRA cells were picked, and the integration of mScarlet was confirmed by PCR followed by Sanger sequencing.

### Generation of cells with doxycycline-inducible TLR4 expression

To generate CEBRA cells that stably express proteins of interest, the PiggyBac transposon system was used^75^. Therefore, the respective sequences encoding the proteins of interest were cloned into the pB-TRE-rtTA donor plasmid, which contains the reverse tetracycline transactivator (rtTA), the tetracycline response element (TRE), and a hygromycin resistance cassette for selection. Point mutations were introduced using site-directed mutagenesis. Cells were electroporated as described above using pTwist_CMVbg_mCherry_T2A_FLAG-hypPB_Neo (encoding hyperactive PiggyBac transposase) and the respective pB-TRE donor plasmid (5 µg total DNA, 1:4). After electroporation, cells were rested for 48 h, followed by selection with hygromycin (500 µg/mL) for two weeks to enrich for cells carrying the integrated construct. Medium was exchanged every 2 – 3 days. Successful stable integration was validated by immunoblotting after doxycycline induction for 24 h.

## Method details

### Clustering eukaryotic TIR-domain proteins

All eukaryotic proteins annotated with Pfam accessions corresponding to TIR domains (PF01582, PF08937, PF10137, PF13676, and PF18567)^76^ as well as proteins annotated with the InterPro accessions IPR000157 and IPR035897, were downloaded from the InterPro database^77^ release 98.0 in January 2024. In addition, a set of 26 known TIR domain structures from bacteria, plants, and animals (Table S7) was used to query the AlphaFold Database UniProt50 (AFDB50)^78,79^ using Foldseek^61^ version 5.53465f0, and hits with probability above 0.95 were retained. The resulting hits were mapped to UniRef50^80^ to collect all cluster members, and only eukaryotic sequences were retained. After merging all sources and removing duplicate IDs, the initial dataset comprised 70,638 sequences. Taxonomic classifications for all proteins were retrieved from UniProt^31^ (release 2024_01). All sequences were filtered for redundancy using the “linclust” option in MMseqs2^58^ (release 13-45111-2021) with the parameter “--min-seq-id 0.9 -c 1”, resulting in 59,752 non-redundant proteins. This dataset was further clustered using the “cluster” option in MMseqs2 with the parameters “--min-seq-id 0.6 -c 0.8”, yielding 19,579 clusters of full-length proteins predicted to encode TIR domains (Table S1). A representative sequence from each cluster was extracted using the “createsubdb” option of MMseqs2. 32 additional well-characterized proteins with TIR domains, which were included in the larger dataset but were not selected as cluster representatives by the “createsubdb” option of MMseqs2, were manually added (Table S2).

### Extraction of TIR domains from proteins

The 3-dimensional structure for each of the cluster representative proteins was predicted using AlphaFold2^60^ with default parameters. Each structure was then searched against a curated set of TIR domain structures^8^ using Foldseek version 5.53465f0^61^. TIR domain boundaries within each protein were defined by identifying the residues corresponding to the median N-terminal and C-terminal positions of hits to the reference TIR domain structures^8^. TIR domain structures were then extracted from the full-length AlphaFold2 predictions using PyMOL^67^ version 3.1.6.1 based on these defined boundaries. This resulted in a total of 16,011 extracted TIR domains (Table S2).

### Structural phylogenetic analysis of TIR domains

To generate a smaller dataset of TIR domains for phylogenetic analysis, the extracted TIR domain sequences were further clustered using the “cluster” option in MMseqs2^58^ with the parameters “--min-seq-id 0.4 -c 0.9”, which resulted in 2,839 clusters, and a representative sequence from each cluster was extracted using the “createsubdb” option of MMseqs2. ‘Singleton’ TIR domain sequences that did not cluster with any other TIR domain, and whose full-length protein sequence did not cluster with any other protein in the previous clustering round, were removed, resulting in a total of 1,490 eukaryotic TIR domains. TIR domains from the 32 well-characterized TIR proteins mentioned above were manually added to the dataset to ensure their inclusion in the subsequent tree.

Structure-guided multiple sequence alignment of TIR domains was performed using FoldMason^62^ version 2.7bd21ed. A phylogenetic tree was constructed using IQ-TREE^63^ version 2.2.0. Selection of the best-fit model of amino acid substitution was inferred using the ModelFinder^81^ function in IQ-TREE, with model VT+F+G4 chosen according to the Bayesian Information Criterion. Node support was computed using 1,000 iterations of the ultrafast bootstrap function^82^ in IQ-TREE (option -bb 1000). The final tree, comprised of 1,522 TIR domains, was visualized with iTOL^64^ v7. Taxonomic clade assignments and the presence of the catalytic glutamic acid residue were determined for each cluster representative TIR domain as described below and used to annotate the corresponding branch on the phylogenetic tree (Table S3).

### Prediction of putative catalytic residues in TIR domains

All extracted TIR domain structures from the cluster representatives were structurally aligned to the TIR domain of ThsB from *Bacillus cereus* MSX-D12^5^, a type I Thoeris protein with experimentally validated catalytic residue E85, using TM-align^65^ version 20170708. For each query TIR domain, the residue whose Cα atom was positioned closest to E85 in the structural alignment was identified. A TIR domain was predicted as putatively catalytic if this aligned residue was a glutamic acid with a Cα atom positioned within 3 Å of the Cα of the E85 residue in ThsB.

### Prediction of additional protein domains

To characterize the domain architecture of TIR-containing proteins, additional domains within these proteins were predicted using HH-suite^59^ version 3.2.0. For each of 16,011 proteins with extracted TIR domains, a multiple sequence alignment (MSA) was constructed using HHblits^83^ against the UniRef30 (2020_06) database with a single search iteration. The resulting MSAs were then searched against the Pfam database using HHsearch^84^. Hits with a probability below 80% or an alignment length below 30 residues were discarded. Overlapping hits (overlap more than 20 residues) were grouped into a single domain.

Domain annotations were then consolidated per protein using Pfam^76^ clan information. In cases in which multiple hits were mapped to the same clan, the clan was used as the representative domain annotation. In cases of overlapping hits with no clan assignment, the most frequently occurring Pfam accession was used. For proteins in which a TIR domain had been previously identified by Foldseek, hits overlapping the TIR domain coordinates were excluded to avoid redundancy with Foldseek-derived TIR annotations. The protein domain annotation was added to Table S2.

### Candidate selection for experimental validation

A subset of eukaryotic TIR domains with predicted catalytic residues was selected for experimental validation, chosen to represent both phylogenetic and structural diversity. For proteins containing additional domains beyond the TIR domain, only the TIR domain sequence was ordered for synthesis; for proteins consisting exclusively of a TIR domain, the full-length sequence was used (Table S4). Structural models of the selected candidates were generated using AlphaFold3^54^ with default parameters. The following SMILES codes were used for the ligands:

cADPR: C1[C@@H]2[C@H]([C@H]([C@@H](O2)N3C=NC4=C3N=CN(C4=N)[C@H]5[C@@H]([C@@H]([C@H]( O5)COP(=O)(OP(=O)(O1)O)O)O)O)O)O.

TAK-242: Fc1cc(c(cc1)N[S](=O)(=O)[C@@H]2CCCC=C2C(=O)OCC)Cl

Structural figures were prepared in PyMOL^67^ version 3.1.6.1 (Figures 2A, S1, S5A, S10, S18).

### Cell lysate preparation for characterization of TIR-derived molecules

*E. coli* TOP10 cells carrying a plasmid expressing the TIR under the control of IPTG promoter were grown at 37 °C with shaking at 200 rpm until reaching an OD_600_ of 0.3 in 50 mL MMB medium. Protein expression was induced with 1mM IPTG and grown for another one or four hours, as indicated in Table S4. Cells were centrifuged at 4 °C, 4,000 *g* for 15 min, and the pellet was frozen in dry ice with ethanol and stored at −80 °C.

To extract cell metabolites, 600 μL of 20 mM potassium phosphate buffer (pH 6) was added to each pellet. Samples were transferred to FastPrep Lysing Matrix B tubes (2 mL; MP Biomedicals, cat. #116911100) and lysed at 4 °C using a FastPrep bead beater for two rounds of 40 s. Tubes were then centrifuged at 4 °C for 10 min at 15,000 *g*. The supernatant was transferred to an Amicon Ultra-0.5 3 kDa centrifugal filter unit (Merck Millipore, cat. no. UFC500396) and centrifuged for 45 min at 4 °C, 15,000 *g*.

### LC-MS analysis of filtered lysates

The sample solutions were analyzed by UPLC–HRMS without dilution. The analyses were carried out on a Waters SYNAPT-XS Q-Tof mass spectrometer (Manchester, UK) with an electrospray ionization (ESI) source. The spectra were recorded in the negative ionization mode within a mass range from 100 to 1200 m/z. The parameters were set as follows: capillary voltage at 1.5 kV, cone gas flow at 50 L/h, source temperature at 140°C, and cone voltage at 20 V (−). The desolvation temperature was set at 600°C, and the desolvation gas (N2) flow rate was set at 800 L/h.

Lock spray was acquired with Leucine Encephalin (m/z=554.2615 in negative mode) at a concentration of 200 ng/mL and a flow rate of 10 μL/min once every 10 s for a 1 s period to ensure mass accuracy. Waters MassLynx v4.2 software was used for data acquisition and data processing.

Standard molecules (2ʹcADPR, 3ʹcADPR, cADPR, His-ADPR, pRib-AMP) were provided by BIOLOG and analyzed in the same conditions as the references. Signals corresponding to the m/z of the cADPR isomers (540.053), His-ADPR (695.126), and pRib-AMP (558.062) were extracted with ±5 ppm mass accuracy, and their retention times were compared with those of the standards. Peaks were reported only when detected in all three repeats, and one representative chromatogram is shown for each sample.

The analytes were separated using the Waters Premier Acquity UPLC system. The gradient elution was achieved with a Waters Acquity Premier HSS T3 Column, 1.8 μm, 2.1 × 100 mm at 0.4 mL/min flow rate, 35°C. Mobile phase A consisted of 20 mM aqueous Ammonium Acetate (Sigma-Aldrich 09691) and mobile phase B consisted of 20 mM Ammonium Acetate in acetonitrile:water ratio of 75:25. The gradient was as follows: 0–2.5 min 0% of B, 2.5-6 min 10% of B.

### Liquid culture growth and toxicity experiments with type VII Thoeris

Overnight bacterial cultures were diluted 1:50 into MMB medium and grown until optical density at 600 nm (OD_600_) of 0.3. Expression was then induced or not with 1 mM IPTG and 800 nM anhydrotetracycline, and grown in 200 µL volumes in 96-well plates at 25 °C. OD_600_ was measured every 10 minutes using a Tecan Infinite 200 plate reader.

#### NAD^+^ detection assay

For cell lysate preparation, cells were grown overnight at 37 °C with 1 mM IPTG induction in a deep 96-well plate in 1 mL MMB media supplemented with ampicillin. Grown cells were harvested by centrifugation at 4,000 *g* for 15 min at 4 °C, and pellets were resuspended in 50 µL of 20 mg/mL lysozyme solution diluted in 100 mM sodium phosphate buffer (pH 7.4). Cells were lysed by two freeze-thaw cycles, each consisting of 15 minutes at room temperature followed by 15 minutes at -80 °C. Cell debris was removed by centrifugation at 4,000 *g* for 15 min at 4 °C, and the supernatant was used for downstream reactions.

For the NAD^+^ detection assay, samples were diluted 1:100 in 100 mM sodium phosphate buffer (pH 7.5). Then, 5 µL of the samples were mixed with 5 µL of NAD/NADH-Glo™ detection reagent (Promega corp.) in a Nunc® 384-well polystyrene plate (Sigma p5991-1CS). Plates were incubated at 25 °C in a TECAN Infinite 200 plate reader, and luminescence was measured every 10 min.

### ThsA NADase assay

The ThsA protein was expressed and purified as previously described^5^, and ThsA NADase assay was performed as previously described^6,85^, with minor modifications. Purified ThsA protein (5 µL, final concentration 100 nM) was added to either 5 µL of cell lysates or 100 mM sodium phosphate buffer (pH 7.5) in black 96-well half-area plates (Corning, cat #3694) to a final reaction volume of 50 µL. Reactions were initiated by adding 5 µL of 1,N6-ethenoadenine dinucleotide (εNAD; Sigma, cat #N2630) to a final concentration of 0.5 mM, immediately before the measurement. Plates were incubated at 25 °C in a Tecan Infinite M200 plate reader, and fluorescence was monitored at 300 nm excitation and 410 nm emission. Reaction rates were calculated from the linear part of the initial reaction.

### ThsE protease reporter assay

ThsE caspase-like protease, synthetic EEKAR–aminomethylcoumarin (AMC) conjugated peptides, and N7-cADPR were produced as described previously^9^. For cell lysate preparation, cells were grown overnight at 37 °C in a deep 96-well plate in 1 mL MMB media supplemented with ampicillin. Overnight cultures were diluted 1:100 into fresh MMB media supplemented with ampicillin and grown for 4 hours at 37 °C. Grown cells were harvested by centrifugation at 4,000 *g* for 15 min at 4 °C, and pellets were resuspended in 50 µL of 20 mg/mL lysozyme solution diluted in 20 mM potassium phosphate buffer (pH 6). Cells were lysed by two freeze-thaw cycles, each consisting of 15 minutes on ice followed by 15 minutes at -80 °C. Cell debris was removed by centrifugation at 4,000 *g* for 15 min at 4 °C, and the supernatant was used for downstream reactions.

ThsE protease activity was measured in a black 384-well plate (Corning 4511) in a total volume of 10 µL containing 3 µL of cell lysate, 1 µM ThsE-containing lysate, 2 µL of 100 µM AMC-conjugated EEKAR peptide, and 4 µL of 100 mM sodium phosphate buffer (pH 7.4). AMC fluorescence (excitation/emission: 341/441 nm) was measured on a Tecan Infinite M200 plate reader at 37 °C every 2 minutes and quantified relative to a 10 µM AMC standard solution.

### Protein purification

TIR domain from *C. reinhardtii* (UniProt ID A0A2K3D750) and its catalytic mutant were cloned into the pET28 expression vector (TWIST Bioscience, USA), which encodes an N-terminal 14xHis tag. Plasmids were transformed into *E. coli* LOBSTR-BL21(DE3)-RIL cells (Kerafast, USA). Cells were grown in 50 mL of MMB media supplemented with kanamycin at 37 °C until mid-log phase (OD_600_∼0.6), then IPTG was added to a concentration of 0.5 mM, and cells were further grown overnight at 16°C, centrifuged, and flash-frozen in dry ice and ethanol. His-tagged protein was purified using 50 µL NEBExpress® Ni-NTA Magnetic Beads (NEB) according to the manufacturerʹs protocol. Elution was done in 100 µL of 500 mM imidazole in IMAC buffer (0.02 M sodium phosphate, 0.3 M NaCl, pH 7.4). The proteins were stored at −80 °C.

#### Enzymatic activity of *C. reinhardtii* TIR domain

100 µM of NAD^+^ was incubated with 1 µM of a purified *C. reinhardtii* TIR in 100 mM sodium phosphate buffer (pH 7.5) for 16 hours at 25 °C. Prior to LC-MS analysis, the products of the reactions were diluted 1:4 in water and were transferred to an Amicon Ultra-0.5 Centrifugal Filter Unit 3 kDa (Merck Millipore, no. UFC500396) and centrifuged for 45 min at 4 °C, 15,000 *g*. LC-MS analysis was performed in the same way as for filtered lysates.

### Multiple sequence alignment of TLR4 and Chlamydomonas TIR homologs

To assess the conservation of the catalytic glutamic acid residue across TLR4 orthologs, all protein sequences were retrieved from the NCBI computed orthologs page^86^ for human TLR4 (Gene ID: 7099) and manually inspected. One ortholog in which the TIR domain was truncated was excluded. The multiple sequence alignment of TLR4 proteins presented in Figure 2B was performed using MAFFT^66^ v7.520 with default parameters and visualized in Jalview^68^ version 2.11.5.1.

To identify homologs of the Chlamydomonas TIR protein, the full-length protein sequence was used as a query in a BLASTp^69^ search against the NCBI non-redundant protein database with default parameters. A set of homologs, selected to represent proteins from different taxonomic groups, was aligned using MAFFT^66^ v7.520 with default parameters and visualized in Jalview^68^ version 2.11.5.1.

### Construction of a tree that includes both eukaryotic and bacterial Thoeris TIRs

TIR proteins from putative Thoeris systems were taken from a previous study^8^. For each TIR-containing gene in the putative Thoeris systems, a structural model was generated using AlphaFold2^60^ with default parameters. TIR domain boundaries were identified, and domains were extracted from these structures, using the same Foldseek-based pipeline described above, resulting in 10,176 TIR domains (Table S5). The catalytic glutamic acid prediction was then applied to the extracted bacterial TIR domains as described above.

The bacterial TIR domain sequences were clustered independently from the eukaryotic sequences using the same MMseqs2 parameters described above. Representative sequences from the bacterial TIR set, supplemented with several experimentally characterized bacterial TIR proteins, were then combined with the set of 1,522 eukaryotic TIRs (Table S3). The combined dataset was aligned using FoldMason^62^ version 2.7bd21ed, and a phylogenetic tree was constructed using IQ-TREE^63^ version 2.2.0 with model VT+G4, selected according to the Bayesian Information Criterion using ModelFinder^81^. Node support was computed using 1,000 ultrafast bootstrap iterations (option -bb 1000)^82^. The final tree was visualized with iTOL^64^ v7 (Figure S7).

### Phylogenetic analysis of Chlamydomonas TIR homologs

To examine the phylogenetic relationships of the *C. reinhardtii* TIR domain and its eukaryotic homologs in the context of bacterial Thoeris TIR domains, sequences from the clade containing the *C. reinhardtii* TIR were extracted from the combined eukaryotic and bacterial TIR domain phylogenetic tree in Figure S7. To produce a more accurate sub-tree, these sequences were aligned using MAFFT^66^ v7.520 with default parameters, and a sub-tree was constructed using IQ-TREE^63^ version 2.2.0. The best-fit substitution model was selected using ModelFinder^81^, with model Q.pfam+G4 chosen according to the Bayesian Information Criterion. Node support was computed using 1,000 iterations of the ultrafast bootstrap function in IQ-TREE (option -bb 1000)^82^. Annotations of putative associated effectors for bacterial systems were obtained from a previously described analysis of Thoeris system predictions^8^. The tree was visualized using iTOL^64^ v7. The list of sequence IDs included in this analysis is provided in Table S6.

### Immunoblotting

Samples were lysed either directly in 2x Laemmli buffer (20 mM Tris pH 6.8, 3.1% DTT (w/v), 4% SDS (w/v), 16% glycerol (v/v), 0.02% bromophenol blue (w/v)) and denatured for 10 min at 95 °C. Otherwise, cells were lysed in RIPA buffer (150 mM NaCl, 50 mM Tris-HCl (pH 7.3), 1% NP-40, 0.1% SDS, 1 mM EDTA) supplemented with cOmplete protease inhibitor cocktail for 15 min on ice and centrifuged for 15 min at 16,000 × g. The supernatant was collected, protein concentration was quantified with a BCA kit, mixed with 6x Laemmli (60 mM Tris pH 6.8, 9.3% DTT (w/v), 12% SDS (w/v), 47% glycerol (v/v), 0.06% bromophenol blue (w/v)), and boiled for 10 min at 95 °C. Proteins were separated by Tris glycine SDS-PAGE and transferred onto ethanol-activated 0.45 µm PVDF membrane at 100 V for 70 min. Membranes were blocked in 3% BSA in TBS-T for 1 h at room temperature, then incubated with the indicated primary and corresponding secondary antibodies. Chemiluminescent signals were acquired using the Immobilon® Forte Western HRP-substrate on the Fusion FX by Vilber Lourmat.

#### ELISA

hIL-6 and hIP-10 ELISAs were conducted according to the supplier’s protocol.

### RT-qPCR analysis

For each condition, cells from four 96-wells were pooled. RNA was isolated using the Genaxxon Total RNA Purification Kit – Tissue following the supplier’s instructions. Next, RNA samples were treated with DNase, and cDNA was generated using the iScript™ cDNA Synthesis Kit (Bio-Rad) following the supplier’s instructions. qPCR was performed using the Takyon® No Rox SYBR®MasterMix dTTP Blue (Eurogentec) according to the manufacturer’s protocol (sequences provided in Supplementary Table x). Data were recorded on the CFX384 Real-Time system (Bio-Rad) and analyzed using CFX Manager v3.1 (Bio-Rad). Primers were designed using the IDT Realtime PCR tool. Recorded *C*_t_ values were normalized to the recorded *C*_t_ of *GAPDH*, and data were plotted as 2(-D*C*_t_) (relative expression).

### Coimmunoprecipitation

For immunoprecipitation of FLAG-tagged endogenous MyD88, CEBRA cells were differentiated in tissue culture-treated 10cm-dishes (600,000 cells/mL) with double the cytokine concentration (M-CSF: 20 ng/mL; IL-3: 20 ng/mL; b-estradiol: 200 nM) for 5 days. Subsequently, cells were stimulated with 200 ng/mL LPS-EB ultrapure for 30 min, washed with ice-cold PBS, and scraped from the plates. For one condition, cells from three dishes were pooled. Whole cell lysates were prepared by adding 1 mL lysis buffer (150 mM NaCl, 50 mM Tris-HCl (pH 7.5), 1% NP-40, 10% glycerol) containing protease/phosphatase inhibitors and incubated on ice for 15 min. Lysates were pre-cleared at 3,000 × g for 5 min at 4 °C, followed by centrifugation at 16,000 × g for 15 min at 4 °C. Whole cell extracts (Input) were quantified with a BCA kit and 1 – 2 mg of this fraction were immunoprecipitated for 4 h at 4 °C using Pierce Anti-DYKDDDDK Magnetic Agarose beads (Thermo Fisher Scientific). After incubation, a sample from the flow-through (FT) was taken. The beads were washed 4 times with lysis buffer. Immunoprecipitated proteins were eluted by incubation in 2x Laemmli for 15 min at 95 °C. Samples were further analyzed by Western blotting.

### Flow cytometry analysis

Cells were detached with ice-cold 2 mM EDTA in PBS for 20 min at 4 °C and washed with PBS. Subsequently, cells were analyzed on a LSRFortessa flow cytometer using FACSDiva v8.0.1 software (BD Biosciences). mScarlet fluorescence was excited using a 561 nm laser and detected using a 585/15 nm bandpass filter. Data were processed using FlowJo v10.9.0. The proportion of mScarlet-positive cells was determined by sequential gating cells on forward scatter (FSC) versus side scatter (SSC), followed by singlet selection using FSC-A versus FSC-H. The mScarlet-positive gate was set according to the unstimulated WT control.

### Live cell analysis

Differentiated CEBRA IL1B^mScarlet^ reporter cells (TRE-CD16 TLR4^WT^ and TRE-CD16-TLR4^E750Q^) were seeded at 100,000 cells/well and stimulated with indicated doxycycline concentrations or R848 (1 µg/mL) for 24 h. Two wells per condition were analyzed, with five images acquired per well at 10× magnification using an Incucyte® SX1 Live-Cell Analysis System and analyzed using the Incucyte® software (GUI version 2025A; Sartorius). Red fluorescent objects were segmented using Top-Hat background subtraction with a radius of 100 µm and a threshold of 2 RCU. Red fluorescence was quantified at the experimental endpoint (24 h) as the mean integrated intensity per detected red object (Red Integrated Intensity Object Mean).

### Quantification and statistical analysis

The average of three triplicates is shown throughout with individual points overlaid, unless stated otherwise. GraphPad Prism 11 was used for data presentation and statistical analysis in Figures 4 and 5. The sample sizes (n) and statistical methods, where applicable, are detailed in the respective figure legends.

## Supplementary Tables

**Supplementary table 1.** Eukaryotic TIR-containing proteins from the UniProt database. Related to Figure 1.

**Supplementary table 2.** Clustered TIR domain-containing proteins with their TIR domain extracted. Related to Figure 1.

**Supplementary table 3.** TIR domains used for the eukaryotic phylogenetic tree. Related to Figure 1.

**Supplementary table 4**. Eukaryotic TIR domains selected for experimental characterization. Related to Figures 2-3.

**Supplementary table 5.** Bacterial TIR-containing proteins predicted to be part of Thoeris systems. Related to figure S7.

**Supplementary table 6.** TIR-containing proteins phylogenetically related to *C. reinhardtii* TIR. Related to Figure S5.

**Supplementary table 7.** A set of known TIR domain structures used to query the AFDB50. Related to Figure 1.

**Supplementary table 8.** Primers used in this study, related to STAR methods.

